# Gastric metabolomics analysis supports *H. pylori*’s catabolism of organic and amino acids in both the corpus and antrum

**DOI:** 10.1101/2020.07.01.183533

**Authors:** Daniela Keilberg, Nina Steele, Sili Fan, Christina Yang, Yana Zavros, Karen M. Ottemann

## Abstract

*Helicobacter pylori* is a chronic bacterial pathogen that thrives in several regions of the stomach, causing inflammation that can vary by site and result in distinct disease outcomes. It is not known, however, whether the host-derived metabolites differ between the two main regions, the corpus and antrum. We thus characterized the metabolomes of mouse gastric corpus and antrum organoids and tissue. The secreted organoid metabolites differed significantly between the corpus and antrum in only seven metabolites: lactic acid, malic acid, phosphoethanolamine, alanine, uridine, glycerol, and isoleucine, with only lactic acid and phosphoethanolamine exceeding two-fold differences. Multiple chemicals were depleted upon *H. pylori* infection including urea, cholesterol, glutamine, fumaric acid, lactic acid, citric acid, malic acid, and multiple non-essential amino acids. Many are known *H. pylori* nutrients and did not differ between the two regions. When tissue was examined, there was little overlap with the organoid metabolites. The two tissue regions were distinct, however, including in antrum-elevated 5-methoxytryptamine, lactic acid, and caprylic acid, and corpus-elevated phospholipid products. Over an 8 month infection time course, the corpus and antrum remained distinct. The antrum displayed no significant changes, in contrast to the corpus which exhibited metabolite changes indicating stress, tissue damage, and depletion of key nutrients such as glutamine and fructose-6-phosphate. Overall, our results suggest the *H. pylori* preferentially uses carboxylic acids and amino acids in complex environments, and these are found in both the corpus and antrum.

## Introduction

The mammalian body is colonized by a rich diversity of bacteria that can utilize a range of nutrients. To establish a long-term colonization, bacteria must sense and adapt to nutrient conditions that likely vary temporally and spatially. One bacterium that can establish a chronic infection is *Helicobacter pylori*, which colonizes two main stomach regions, the corpus, also called the fundus in mice, and the antrum. Little is known about the nutrients *H. pylori* utilizes in these two regions and how they vary over space and time.

*H. pylori* colonizes the stomachs of many people: 50% worldwide and 35% in the United States (1). The infection is acquired in childhood, and becomes chronic and persistent unless treated with antibiotics. Approximately 85% of infected people are asymptomatic, developing only a low-level inflammation. However, around 15% of infected people will develop serious *H. pylori* triggered disease during their life, of either a gastric or duodenal ulcer or gastric cancer (2, 3).

The corpus and antrum are distinct from each other in several ways. In each of these regions, the epithelium is highly invaginated into gastric glands that each contain stem cells (4, 5). These stem cells give rise to a distinct set of epithelial cells in each area. Both areas contain cells that secrete mucus apically, but the corpus is specialized for digestive functions while the antrum is specialized for endocrine ones (6). The corpus digestive functions are created via acid-secreting parietal cells along with pepsinogen- and lipase-secreting chief/zymogenic cells, all secreted apically. The antrum endocrine function is created by gastrin-producing G cells, which is secreted basolaterally. Overall, these different cell types create a distinct milieu in each region, which presumably requires distinct *H. pylori* abilities for colonization. There is some support for this idea: *H. pylori* mutants lacking chemotaxis have a greater colonization defect in the antrum as compared with the corpus (7–9), while mutants lacking the cytotoxin-associated pathogenicity island (*ca*gPAI) genes *cagA* or *cagY* have a greater colonization defect in the corpus as compared to the antrum (10). However, the nature of the differences that *H. pylori* encounters in these two regions is not known.

Several studies have compared *H. pylori* colonization dynamics between the mouse corpus and antrum (8, 9). *H. pylori* infections begin corpus dominant, by 5-10-fold higher amounts compared with the antrum. Over the first two months, however, *H. pylori* multiplies to a great extent in the antrum, such that the infection becomes antral dominant from one week post infection to ∼ 2 months post infection (8, 9). After two months of infection, the *H. pylori* numbers decline more-so in the antrum and the infection becomes corpus dominant again. These experiments suggest the antrum switches between a challenging and favorable environment for *H. pylori* growth. In agreement with this idea, bacterial chemotaxis, the ability to sense environmental conditions and move toward beneficial ones and away from harmful ones, is particularly critical in the antrum (7, 9). Overall, these colonization patterns suggest that host metabolites might differ between the corpus and the antrum, and possibly change over the course of the infection.

Knowledge about the corpus and antrum conditions is important because *H. pylori*-triggered diseases differ in these two areas. Inflammation localized to the antrum leads mainly to ulcers and few cancers, inflammation in the corpus correlates with intestinal-type gastric cancer, and inflammation throughout the stomach increases the risk of diffuse-type gastric cancer (3, 11, 12). It is not known whether there are metabolites differences between these regions. Metabolomics studies of gastric tissue have been limited. The total mouse gastric metabolome was analyzed over an *H. pylori* infection time course, but the separate corpus and antrum metabolites were not characterized (13). They did find evidence of upregulation of glycolysis, TCA and choline pathway intermediates, but the findings were not consistent across time points. Other studies have compared the metabolomes of human gastric cancerous and normal tissue (Reviewed in (14)), in part with the idea that these metabolites might be biomarkers of disease (15). Those studies were noted to have limitations, but *in toto* suggested that cancerous tissue has elevated amounts lactic acid, citric acid, fumaric acid, glutamine, glutamate, and valine as well as several free fatty acids. To help fill gaps in our knowledge, we thus embarked on a study of the corpus and antrum metabolites, to describe them and evaluate how *H. pylori* infection affected them. We report here that a handful of metabolites differ between these compartments. Several are depleted upon *H. pylori* infection, and thus may represent preferred nutrient sources for *H. pylori*. We additionally find that the antrum is surprisingly stable pre- and post-infection, while the corpus undergoes substantial alterations. Overall, our results are consistent with *H. pylori* preferential catabolism of carboxylic acids and amino acids in both the corpus and the antrum.

## Results

### Secreted metabolites of antrum and corpus gastric organoids are similar

We investigated the differences in corpus/fundus and antrum first using a mouse organoid system, because this approach would allow us to isolate secreted metabolites that would be accessible to *H. pylori*. For this experiment, mouse gastric glands were isolated from the corpus or antrum, cultured as spherical organoids. This culture method results in representation of all standard stomach epithelial cells, including surface pit, mucous neck, chief, endocrine and parietal cells (16). After development of spheroids, the organoids were transferred to collagen-coated dishes to allow them to flatten out as two-dimensional (2D) organoids. For the 2D organoid culture, the medium is very rich and contains tissue culture medium (DMEM), fetal bovine serum, and multiple growth factor supplements. For metabolomics analysis, the culture supernatant was collected, filtered to remove cells, and analyzed using gas chromatography time of flight (GC-TOF) mass spectroscopy for various metabolites, including carbohydrates, sugar phosphates, amino acids, hydroxyl acids, free fatty acids, purines, and pyrimidines.

Using this approach, a total of 352 compounds were found to be secreted by the organoids, of which 132 (38%) were identified chemicals, while the other 220 (62%) were unknown chemical structures (Supplemental Table 1). Principal component analysis suggested that the organoids differed only minimally from the media (Fig. 1A), which was consistent with the richness of the organoid media. Specifically, the corpus organoids had no significant differences from media, while the antrum organoids differed significantly in 13 compounds, of which 12 were identified chemicals (Supplemental Table 2). Nine chemicals were decreased in the antral organoids as compared to the media, suggesting they may be used by the antral organoid cells. These nine included the amino acids aspartic acid, leucine, serine, isoleucine, phenylalanine, and valine, and the chemicals glycerol, ethanolamine and nicotinamide (Supplemental Table 2).Three metabolites, however, were elevated 1.7-3.9 fold by the antral organoid system, suggesting they may be excreted by the cells: the carboxylic acids lactic acid and malic acid, and the phospholipid component phosphoethanolamine. This analysis thus suggests that media collected after exposure to corpus organoids did not change much in composition, while exposure to antral organoids lead to a significant increase of lactic acid, malic acid and phosphoethanolamine.

**Fig. 1.**
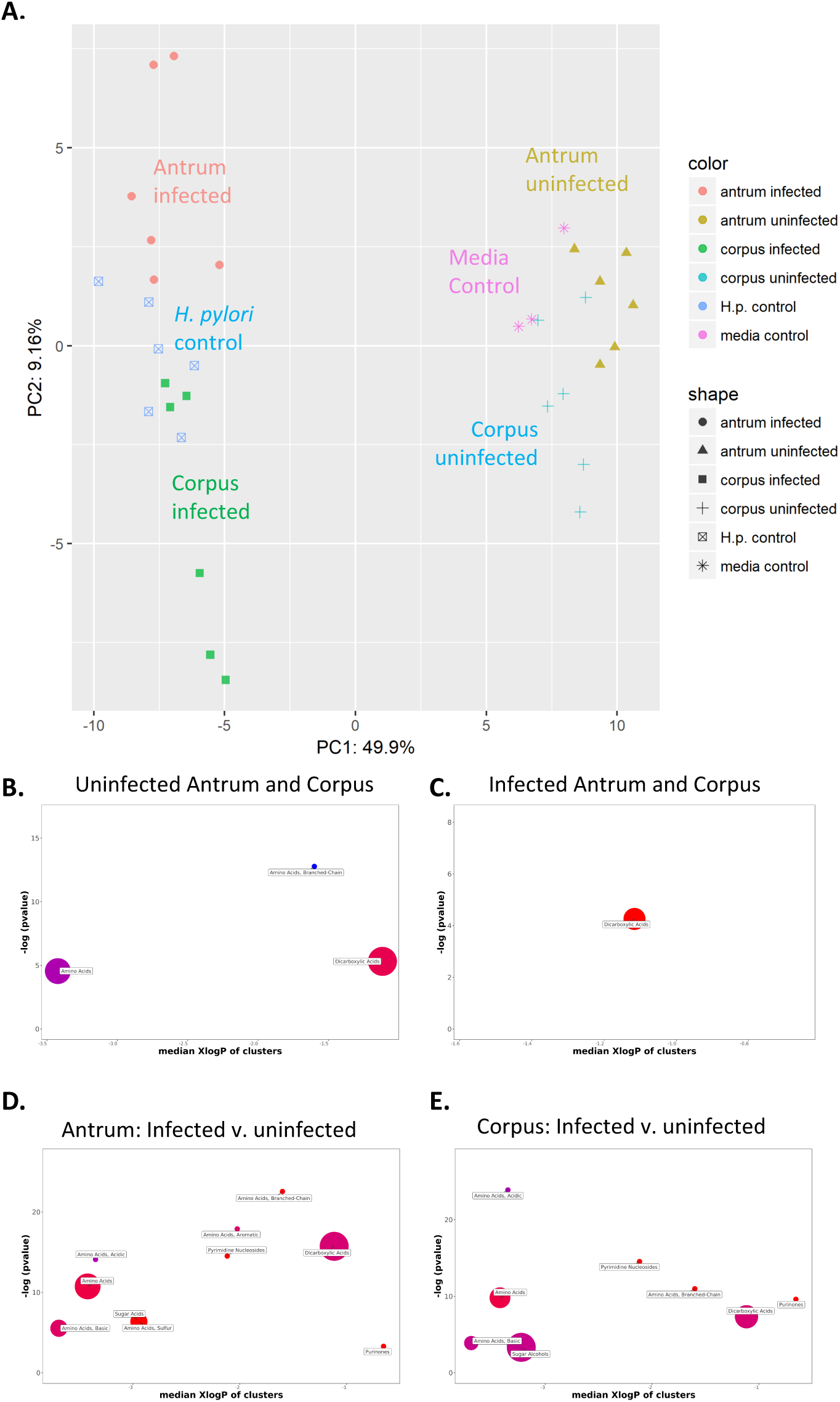
Comparison of metabolites from 2D organoid culture supernatants of mouse gastric antrum and corpus, with and without infection *by H. pylori* SS1. Control samples include organoid media alone and *H. pylori* alone. A. Principle Component Analysis (PCA). B-E. ChemRICH enrichment analysis for pairwise comparison as indicated. The Y axis shows the most significantly altered clusters at the top, with enrichment p-values given by the Kolmogorov–Smirnov-test. Only enrichment clusters are shown that are significantly different at p<0.05. The X axis shows the clusters similarities to each other. Each node reflects a significantly altered cluster of metabolites. Node sizes represent the total number of metabolites in each cluster set. The node color scale shows the proportion of increased (red), decreased (blue) compounds, or both increased and decreased metabolites (purple). For each sample, N = 6 except media alone where N = 3.

We next compared the antrum and corpus organoids to each other (Fig. 1A). 26 named compounds differed between the two, with seven of these significant (Table 1). Antral organoids were significantly enriched over the corpus organoids for lactic acid, malic acid, and phosphoethanolamine, as well as the amino acid alanine. Corpus organoids, on the other hand, were enriched for the pyrimidine nucleoside uridine, glycerol, and the amino acid isoleucine. For the most part, these differences were not more than 2-fold, but a few compounds were more substantial. The antrum was over 2-fold enriched for lactic acid and phosphoethanolamine; fumaric acid was also highly enriched but did not reach significance (Table 1). ChemRICH chemical similarity analysis (17) using the larger set of non-FDR corrected significant compounds and fold changes identified that the corpus-antrum organoid differences were dominated by differences in general amino acids, branched chain amino acids, and dicarboxylic acids (Fig. 1B and Supplemental Table 3A). These analyses show that corpus and antrum organoids display some differences between each other, dominated by the antral presence of carboxylic acids and phosphoethanolamine.

**Table 1.**
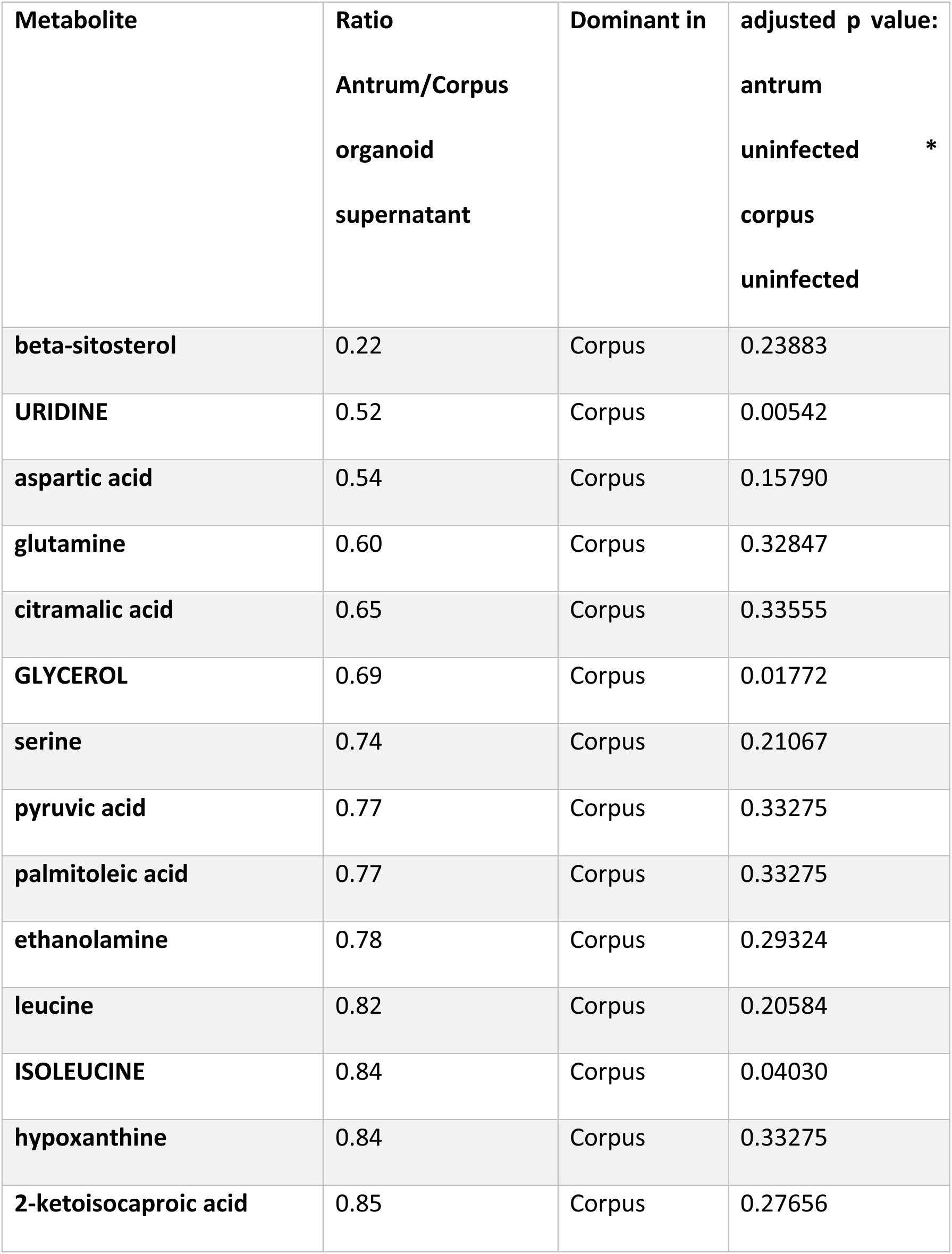
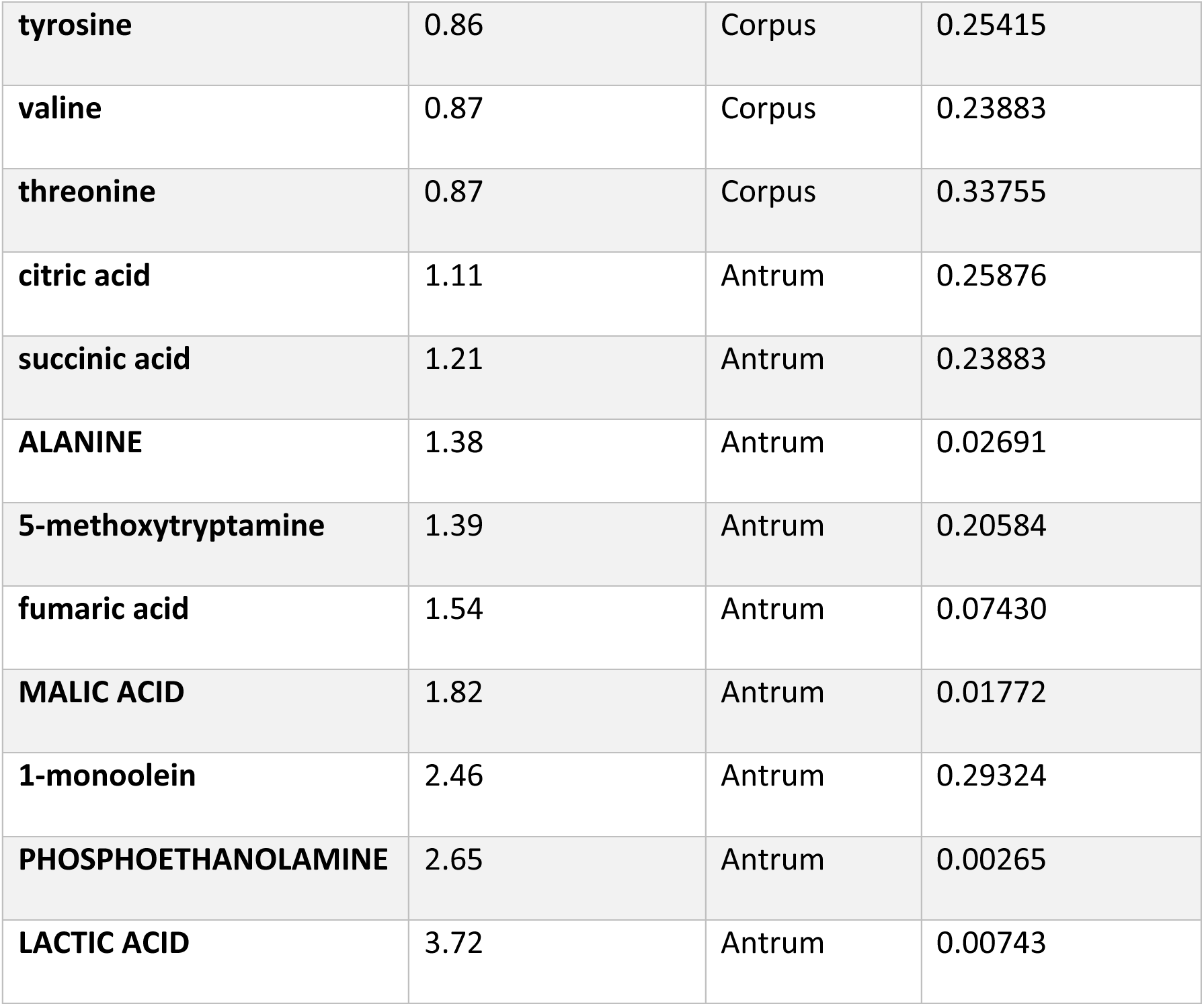
Uninfected gastric organoid supernatant was collected and analyzed for total metabolites. Compounds that differed between corpus and antrum organoids, ranked by fold difference. Red capitol letter text indicates those that are highly significant in the whole data set after applying false discovery parameters, but all compounds that were significant when comparing just these two samples are shown. N= 6 for each group.

| **Metabolite** | **Ratio**<br>**Antrum/Corpus**<br>**organoid**<br>**supernatant** | **Dominant in** | **adjusted p value:**<br>**antrum**<br>**uninfected \***<br>**corpus**<br>**uninfected** |
| --- | --- | --- | --- |
| **beta-sitosterol** | 0.22 | Corpus | 0.23883 |
| **URIDINE** | 0.52 | Corpus | 0.00542 |
| **aspartic acid** | 0.54 | Corpus | 0.15790 |
| **glutamine** | 0.60 | Corpus | 0.32847 |
| **citramalic acid** | 0.65 | Corpus | 0.33555 |
| **GLYCEROL** | 0.69 | Corpus | 0.01772 |
| **serine** | 0.74 | Corpus | 0.21067 |
| **pyruvic acid** | 0.77 | Corpus | 0.33275 |
| **palmitoleic acid** | 0.77 | Corpus | 0.33275 |
| **ethanolamine** | 0.78 | Corpus | 0.29324 |
| **leucine** | 0.82 | Corpus | 0.20584 |
| **ISOLEUCINE** | 0.84 | Corpus | 0.04030 |
| **hypoxanthine** | 0.84 | Corpus | 0.33275 |
| **2-ketoisocaproic acid** | 0.85 | Corpus | 0.27656 |
| **tyrosine** | 0.86 | Corpus | 0.25415 |
| **valine** | 0.87 | Corpus | 0.23883 |
| **threonine** | 0.87 | Corpus | 0.33755 |
| **citric acid** | 1.11 | Antrum | 0.25876 |
| **succinic acid** | 1.21 | Antrum | 0.23883 |
| **ALANINE** | 1.38 | Antrum | 0.02691 |
| **5-methoxytryptamine** | 1.39 | Antrum | 0.20584 |
| **fumaric acid** | 1.54 | Antrum | 0.07430 |
| **MALIC ACID** | 1.82 | Antrum | 0.01772 |
| **1-monoolein** | 2.46 | Antrum | 0.29324 |
| **PHOSPHOETHANOLAMINE** | 2.65 | Antrum | 0.00265 |
| **LACTIC ACID** | 3.72 | Antrum | 0.00743 |

### Antrum and corpus organoid supernatants show a metabolomic response to *H. pylori* infection

We next examined how the antrum and corpus organoid supernatants changed after *H. pylori* infection. Both types of organoids support *H. pylori* colonization (18, 19). After preparation, the 2D organoids were infected by adding the mouse-adapted wild-type *H. pylori* strain SS1 to the media. Infections proceeded for 24 hours, and then the supernatant was collected, filtered, and analyzed as above. Out of 132 identified chemicals, infection resulted in a change of 46 and 51 identified chemicals in the antrum and corpus, respectively (Table 2-3; Supplemental Table 4-5). Changes in both types of organoid supernatants mapped to multiple categories of metabolites, but most substantially to amino acids and dicarboxylic acids, and additionally sugar alcohols in the corpus (Fig. 1D-E, Supplemental Table 3). These changes resulted in a substantial shift in the principal components between uninfected and infected samples (Fig. 1A).

**Table 2.** Compounds that differed significantly between infected and uninfected antral organoid supernatant. Antral organoids were cultured as 2D organoids and infected with wild-type H. pylori SS1 for 24 hours. After infection, the culture supernatant was collected and filtered to remove bacteria. P-values are corrected for multiple comparison problem using Benjamini-Hochberg procedure. N = 6 samples/organoid group and 3 for the *H. pylori* and media controls. ** indicates the only increased compound that was different from the *H. pylori* alone samples.

| **Metabolite** | **Ratio**<br>**Infected/Uninfected** | **FDR corrected P**<br>**value** |
| --- | --- | --- |
| **urea** | 0.04 | 0.00000 |
| **glutamine** | 0.10 | 0.00010 |
| **malic acid** | 0.23 | 0.00001 |
| **n-acetylglutamate** | 0.23 | 0.00036 |
| **serine** | 0.34 | 0.00007 |
| **fumaric acid** | 0.36 | 0.00010 |
| **lactic acid** | 0.40 | 0.05266 |
| **citric acid** | 0.44 | 0.00000 |
| **cholesterol** | 0.51 | 0.00612 |
| **aspartic acid** | 0.65 | 0.03809 |
| **oxoproline** | 0.74 | 0.03809 |
| **tyrosine** | 0.78 | 0.01238 |
| **isoleucine** | 1.45 | 0.00314 |
| **inosine** | 1.50 | 0.01138 |
| **threonine** | 1.53 | 0.00040 |
| **myristic acid** | 1.55 | 0.03049 |
| **ethanolamine** | 1.59 | 0.00218 |
| **asparagine** | 1.64 | 0.00011 |
| **tryptophan** | 1.67 | 0.00727 |
| **valine** | 1.67 | 0.00091 |
| **ribose** | 1.88 | 0.00001 |
| **5-methoxytryptamine** | 1.89 | 0.00230 |
| **glyceric acid** | 1.89 | 0.01138 |
| **succinic acid** | 2.03 | 0.00001 |
| **pseudo uridine** | 2.05 | 0.03397 |
| **phenylalanine** | 2.07 | 0.00009 |
| **alanine** | 2.09 | 0.00002 |
| **leucine** | 2.11 | 0.00001 |
| **glycine** | 2.26 | 0.00064 |
| **glycerol-alpha-phosphate** | 2.54 | 0.02841 |
| **citramalic acid** | 2.76 | 0.00010 |
| **uracil** | 3.00 | 0.00262 |
| **pyruvic acid** | 3.06 | 0.00010 |
| **thymidine** | 3.13 | 0.00006 |
| **glycerol** | 3.57 | 0.00002 |
| **uric acid** | 4.41 | 0.00006 |
| **homoserine** | 5.02 | 0.00001 |
| **cysteine** | 5.34 | 0.00603 |
| **uridine** | 5.81 | 0.00000 |
| **cystine** | 5.82 | 0.00006 |
| **xanthine** | 7.58 | 0.00225 |
| **lysine** | 9.08 | 0.00120 |
| **glutamate** | 9.68 | 0.00000 |
| **histidine** | 12.79 | 0.00007 |
| **methionine** | 14.41 | 0.01977 |
| **taurine** | 43.87 | 0.00719 |

**Table 3.** Compounds that differed significantly between infected and uninfected corpus organoid supernatant. Corpus organoids were cultured as 2D organoids and infected with wild-type *H. pylori* SS1 for 24 hours. After infection, the culture supernatant was collected and filtered to remove bacteria. P-values are corrected for multiple comparison problem using Benjamini-Hochberg procedure. N = 6 samples/organoid group and 3 for the *H. pylori* and media controls. Note: of the increased compounds, none differed between the infected corpus organoids and *H. pylori* alone.

| **Metabolite** | **Ratio**<br>**Infected/Uninfected** | **FDR**<br>**corrected**<br>**P value** |
| --- | --- | --- |
| **lactic acid** | 0.01 | 0.00000 |
| **urea** | 0.02 | 0.00000 |
| **glutamine** | 0.06 | 0.00000 |
| **aspartic acid** | 0.17 | 0.00001 |
| **lactamide** | 0.27 | 0.00001 |
| **n-acetylglutamate** | 0.31 | 0.00035 |
| **malic acid** | 0.32 | 0.00000 |
| **serine** | 0.36 | 0.00006 |
| **fumaric acid** | 0.39 | 0.00193 |
| **capric acid** | 0.52 | 0.01265 |
| **citric acid** | 0.53 | 0.00007 |
| **tyrosine** | 0.60 | 0.00042 |
| **benzoic acid** | 0.62 | 0.01764 |
| **oxoproline** | 0.65 | 0.00027 |
| **threitol** | 0.81 | 0.02178 |
| **guanosine** | 0.81 | 0.02083 |
| **isoleucine** | 1.17 | 0.02577 |
| **threonine** | 1.21 | 0.02849 |
| **oxalic acid** | 1.22 | 0.04623 |
| **stearic acid** | 1.23 | 0.04208 |
| **inosine** | 1.33 | 0.00723 |
| **1-monostearin** | 1.34 | 0.04585 |
| **valine** | 1.36 | 0.00072 |
| **phenylalanine** | 1.45 | 0.03142 |
| **myristic acid** | 1.48 | 0.01723 |
| **phosphoethanolamine** | 1.51 | 0.00293 |
| **hypoxanthine** | 1.52 | 0.03807 |
| **glycine** | 1.53 | 0.00049 |
| **glycerol-alpha-phosphate** | 1.56 | 0.00738 |
| **glyceric acid** | 1.61 | 0.00098 |
| **pseudo uridine** | 1.61 | 0.00788 |
| **leucine** | 1.65 | 0.00065 |
| **sucrose** | 1.91 | 0.01583 |
| **glycerol** | 1.99 | 0.03715 |
| **5-methoxytryptamine** | 2.00 | 0.00009 |
| **ribose** | 2.01 | 0.00024 |
| **2-hydroxyglutaric acid** | 2.06 | 0.00392 |
| **alanine** | 2.17 | 0.00002 |
| **succinic acid** | 2.19 | 0.00002 |
| **thymidine** | 2.48 | 0.00006 |
| **uric acid** | 2.52 | 0.04075 |
| **uracil** | 2.56 | 0.01117 |
| **homoserine** | 3.39 | 0.00039 |
| **thymine** | 3.81 | 0.00000 |
| **uridine** | 3.94 | 0.00002 |
| **cystine** | 7.75 | 0.00000 |
| **xanthine** | 7.83 | 0.00008 |
| **glutamate** | 11.52 | 0.00000 |
| **taurine** | 11.87 | 0.00077 |
| **lysine** | 12.66 | 0.00000 |
| **histidine** | 16.62 | 0.00000 |

Several of the changed metabolites were depleted with *H. pylori* infection, and these metabolites are generally consistent with what is known about *H. pylori* metabolism. For example, urea was highly depleted in both corpus and antrum systems (Table 2; Fig. 2-3). This decrease agrees with *H. pylori*’s robust urease and its known ability to find urea via chemotaxis (Huang et al. 2015). Similarly, glutamine was highly depleted, consistent with *H. pylori* ’s gamma glutamyl transferase (GGT) which breaks glutamine into glutamate and ammonia (20, 21). Lactic acid was highly depleted in each system too, consistent with *H. pylor*i ’s ability to use lactic acid as a carbon or energy source (22). Additional amino acids or amino acid derivatives were substantially depleted as well: serine, N-acetylglutamate, aspartic acid, oxoproline/pyroglutamic acid, and tyrosine. Only one of these is an essential amino acid—serine, and only for some strains (23). The other amino acids are likely used as *H. pylori* carbon and nitrogen source, as has been shown for aspartic acid (24). Also used up were several additional carboxylic acids including malic acid, fumaric acid, and citric acid.

**Fig. 2.**
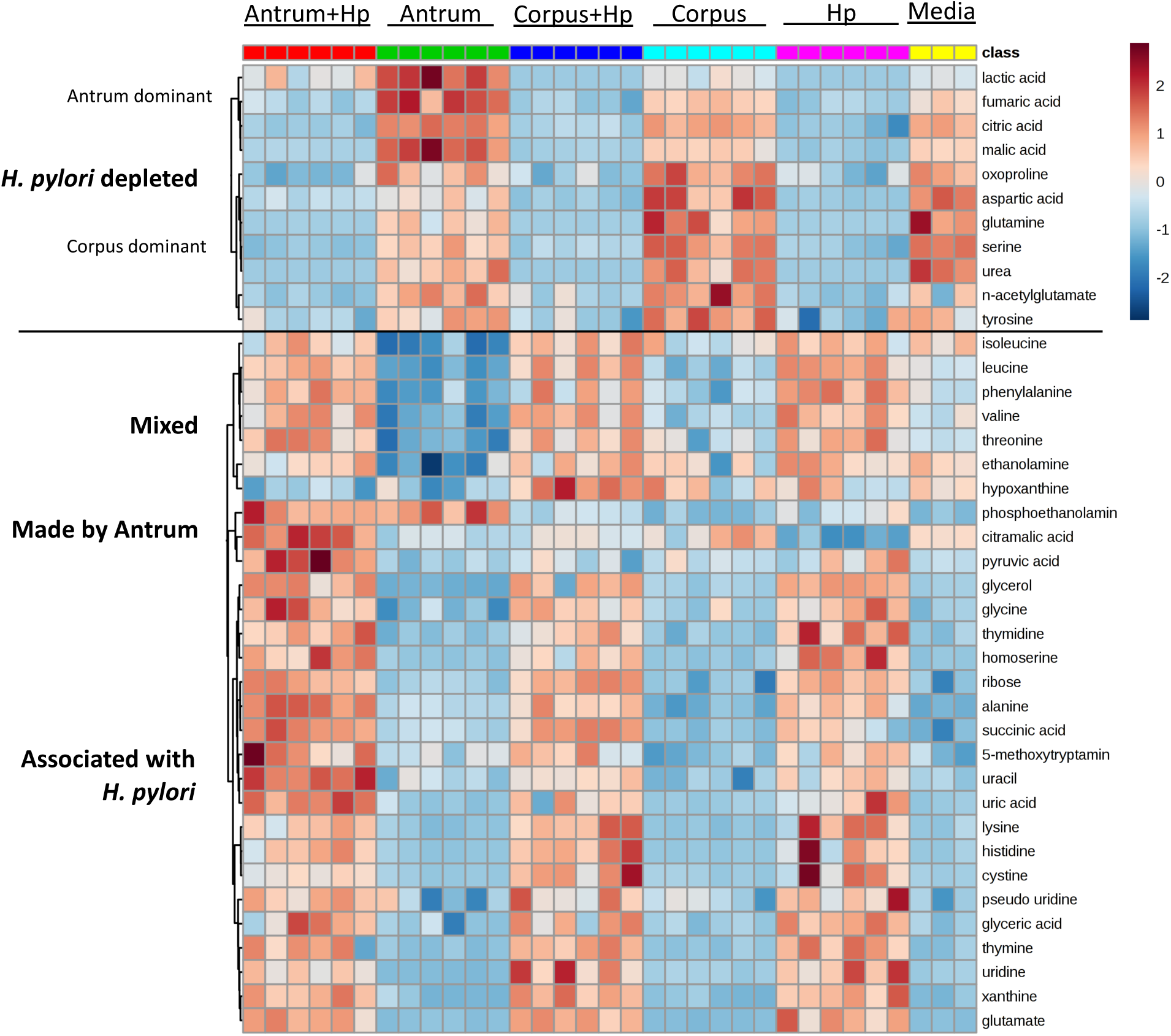
Heat map showing metabolite alterations in organoid supernatant following *H. pylori* infection. Heat map was generated using Metaboanalyst to show the top 40 most significantly changed compounds identified by ANOVA across groups. Left side labels indicate a description of the major nodes.

**Fig. 3.**
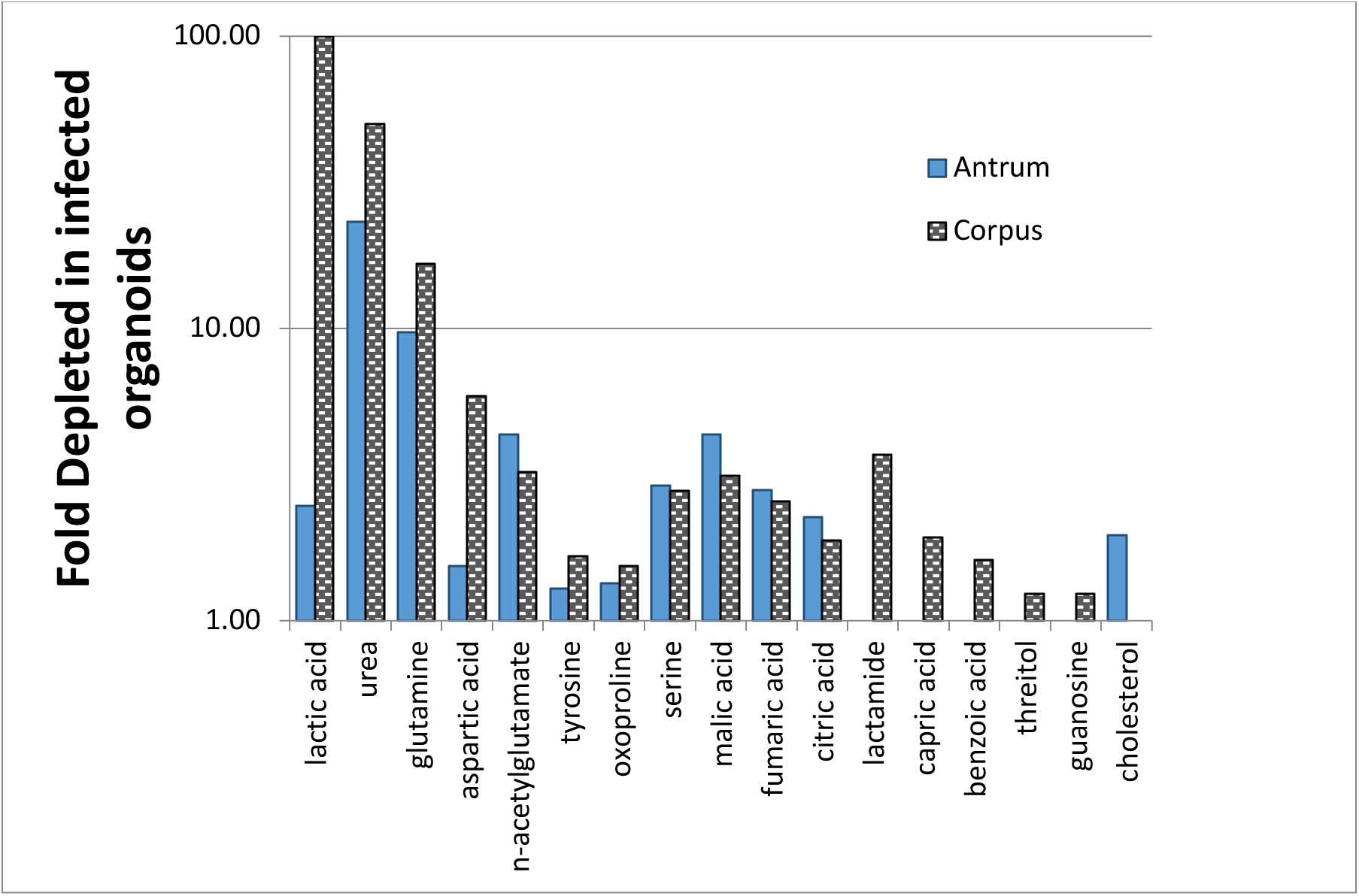
Comparison of supernatant compounds depleted upon infection in corpus and antrum organoids.

Overall, the metabolite usage was similar but non identical between the corpus and antrum organoids, suggesting *H. pylori* has a fairly similar metabolism in both regions. There were some differences, however. For example, cholesterol, a requirement for *H. pylori* growth (25, 26), was substantially depleted in infected antrum organoids but not corpus ones (Fig. 3), although present in both. Additionally, there were several compounds that were lowered more in the corpus, including the carboxamide lactic acid derivative lactamide, the saturated medium chain fatty acid capric acid (also called decanoic acid), benzoic acid, the sugar alcohol threitol, and the purine guanosine. Again, these compounds were present in both organoids, but not highly used in the antrum. Overall, these results suggest that *H. pylori* depletes multiple metabolites in both corpus and antrum organoids that appear to be key nutrients and growth substrates, with some region-specific utilization.

Several compounds were increased in infected organoids as compared to uninfected ones (Table 2, Table 3). The vast majority of these compounds appeared to come from *H. pylori*, as they were also present in bacteria alone samples (Fig. 2). Indeed, when we compared the infected antral organoids to the *H. pylori* samples, only one compound was significantly increased: the malic acid analog citramalic acid (Fig. 2).

After infection, corpus and antrum organoids appeared more distinct than before infection as seen by a shift and separation of the principle components (Fig. 1A), although the number of significantly different metabolites were about the same before and after infection (Table 1, Table 4). After infection, there were five significantly differing compounds that were all found in higher levels in the antrum than in the corpus. The most striking change was in lactic acid, which was heavily used in the corpus and became 150-fold more abundant in the antrum. Phosphoethanolamine also remained enriched in the antrum, staying at about 2-fold over the corpus. The antrum was newly enriched for lactamide, uracil, and citramalic acid (Table 4). Employing ChemRICH enrichment analysis suggested that substantial differences in amino acids were lost, and that differences were dominated by those in dicarboxylic acids (Fig. 1C). Overall, these analyses suggest that *H. pylori* uses a discrete set of organoid-secreted metabolites, dominated by urea, glutamine, lactic acid, fumaric acid, citric acid, and specific amino acids. For the most part, these are used similarly but non-identically between the types of organoids with more utilization in the corpus. This differential resulted in the post-infection metabolomes differing from the pre-infection ones.

**Table 4.** Comparison of supernatant of infected antrum and corpus organoids.

| **Metabolite** | **Fold change antrum**<br>**infected / corpus infected** | **FDR**<br>**corrected**<br>**P value** |
| --- | --- | --- |
| **uracil** | 1.71 | 0.0071 |
| **phosphoethanolamine** | 1.97 | 0.00018 |
| **citramalic acid** | 2.06 | 0.0071 |
| **lactamide** | 5.37 | 0.00599 |
| **lactic acid** | 151.68 | 0.00018 |

### The uninfected mouse tissue differs between antrum and corpus

We next assessed how uninfected mouse gastric tissue compared with the gastric organoids analyzed above. First, mouse gastric tissue had substantially fewer total compounds as compared with organoids: 194 total compounds in the gastric tissue as compared with 352 in the organoids. A somewhat higher percentage of the tissue metabolites were known: 52% in the tissue as compared to 38% in the organoids (Suppl. Table 7). Between the two regions, there were 15 identified compounds that differed significantly between the corpus and the antrum tissue (Table 5), a number that was about 2-fold greater than the number of metabolites that were different between the uninfected antrum and corpus organoids (Table 1). Generally, most of these differing compounds were enriched in the antrum. Specifically, the antrum was enriched for several compounds used by *H. pylori* in the organoids, including lactic acid, glutamine, capric acid, and benzoic acid (Fig. 3), as well as other *H. pylori* nutrients such as pyruvic acid. The corpus was enriched for a few other compounds used up in the organoid system, malic acid and fumaric acid, (Table 5). Overall, the majority of the differing compounds were elevated in the antrum.

**Table. 5.** Uninfected mouse gastric tissue was collected from female C57Bl6/N mice and analyzed for total metabolites. Shown are all compounds that differed when comparing the two groups, with capitol letters indicating differences that were highly significant in the whole data after applying false discovery corrections. ** indicates similarly different in the corpus or antrum organoids. N = six mice for each group.

| **Metabolite** | **Ratio Antrum/Corpus gastric tissue** | **Dominant in** | **FDR correct p value** |
| --- | --- | --- | --- |
| **ARACHIDONIC ACID** | 0.06 | Corpus | 0.00500 |
| **LINOLEIC ACID** | 0.14 | Corpus | 0.00600 |
| **OLEIC ACID** | 0.18 | Corpus | 0.00700 |
| **GLYCEROL-ALPHA-PHOSPHATE** | 0.37 | Corpus | 0.00600 |
| **ETHANOLAMINE\*\*** | 0.42 | Corpus | 0.00100 |
| **adenosine-5-monophosphate** | 0.43 | Corpus | 0.06200 |
| **1-monoolein** | 0.45 | Corpus | 0.05500 |
| **SQUALENE** | 0.66 | Corpus | 0.03300 |
| **fumaric acid** | 0.74 | Corpus | 0.10600 |
| **inosine** | 0.75 | Corpus | 0.06900 |
| **malic acid** | 0.8 | Corpus | 0.10600 |
| **threonine** | 1.24 | Antrum | 0.07600 |
| **phosphate** | 1.29 | Antrum | 0.06200 |
| **creatinine** | 1.43 | Antrum | 0.10700 |
| **palmitic acid** | 1.51 | Antrum | 0.07100 |
| **caprylic acid** | 1.53 | Antrum | 0.10600 |
| **capric acid** | 1.54 | Antrum | 0.08600 |
| **pelargonic acid** | 1.56 | Antrum | 0.08800 |
| **FUCOSE** | 1.61 | Antrum | 0.00800 |
| **benzoic acid** | 1.66 | Antrum | 0.06200 |
| **oxalic acid** | 1.69 | Antrum | 0.06900 |
| **uridine** | 1.73 | Antrum | 0.10700 |
| **MANNITOL** | 1.74 | Antrum | 0.00800 |
| **URIC ACID** | 1.76 | Antrum | 0.01300 |
| **LACTIC ACID\*\*** | 1.79 | Antrum | 0.01200 |
| **glutamine** | 1.83 | Antrum | 0.06000 |
| **succinic acid\*\*** | 1.84 | Antrum | 0.10500 |
| **hydroxylamine** | 2.01 | Antrum | 0.09700 |
| **DODECANOL** | 2.03 | Antrum | 0.04400 |
| **fructose-6-phosphate** | 2.13 | Antrum | 0.06000 |
| **hexose-6-phosphate** | 2.3 | Antrum | 0.07000 |
| **OCTADECANOL** | 2.43 | Antrum | 0.03300 |
| **glucose-6-phosphate** | 2.56 | Antrum | 0.08000 |
| **2-MONOPALMITIN** | 3.5 | Antrum | 0.01100 |
| **glucose** | 4.37 | Antrum | 0.06200 |
| **SUCROSE** | 4.89 | Antrum | 0.01200 |
| **5-METHOXYTRYPTAMINE\*\*** | 6.06 | Antrum | 0.00600 |
| **pyruvic acid** | 6.7 | Antrum | 0.05600 |

When comparing the metabolites of the gastric tissue to the organoids, there was not substantial overlap (Table 1 and Table 5). This lack of similarity is perhaps not surprising, as the mouse tissue sample was whole sample, not restricted to secreted molecules and was cultured in the lab for multiple days. The cultured organoid cells are fed predominantly glucose and glutamine, which may be different than their in vivo metabolism. Therefore, it may be that *H. pylori* is exposed to different metabolites in the two systems.

### The corpus and antrum mouse tissue remain distinct post infection

We then infected mice and evaluated how the metabolomes changed over time. Samples were collected from mice infected for 3 days, 28 days, and 8 months, representing early and chronic infections and confirmed for infection (Supplemental Fig. 1). We then used these samples to see whether the corpus and antrum metabolite differences were stable across this time course. Generally, as the infection proceeded, there were fewer compounds that differed between the two regions. Using the full set of different compounds (not FDR corrected), the differing metabolites decreased from 37, to 14, 25, and 29 at the 3-day, 28-day, and 8-month timepoints respectively (Table 6). Compounds that were consistently distinct between the corpus and antrum included the antrum-elevated 5-methoxytryptamine, lactic acid, and the fatty acid caprylic/octanoic acid. Chemicals that were regularly high in the corpus included multiple products that derive from phospholipids, including arachidonic acid, linoleic acid, oleic acid, and glycerol alpha phosphate (Table 6). Lastly, several compounds were distinct at the outset, before infection, but became more similar between regions after infection, including monoacylglycerol 2-monopalmitin (also called Glyceryl 2-palmitate), the sugars glucose and sucrose, and the carboxylic acids pyruvic acid and succinic acid. Taken together, the data suggests that the corpus and antrum remain distinct throughout the infection, with relatively modest changes, but may become slightly more similar over time.

**Table 6.** Ratios in the mouse antrum and corpus tissue over time post infection. For each time point, all named metabolites that differed using a non-FDR corrected P value (T test, two sample) were used. Orange color indicates up in antrum. Blue indicates down in antrum. Italics indicates one time point.

|  | Ratio Antrum/Corpus |  |  |  |
| --- | --- | --- | --- | --- |
| Metabolite | Uninfected | 3 day | 28 days | 8 months |
| 1-monoolein<br>(Monooleoylglycerol) | 1.93 | 1.07 | 0.43 |  |
| 2-monopalmitin | 3.50 |  |  |  |
| 3-hydroxybutyric acid |  |  |  | 1.89 |
| 3-phosphoglycerate |  |  | 3.03 | 3.71 |
| 5-methoxytryptamine | 6.06 | 4.06 | 5.54 | 3.63 |
| adenosine-5-monophosphate | 0.43 |  | 0.56 |  |
| arachidic acid |  |  | 1.38 |  |
| arachidonic acid | 0.06 | 0.29 | 0.22 | 0.17 |
| aspartic acid |  |  |  | 0.57 |
| benzoic acid | 1.66 |  |  | 1.75 |
| capric acid | 1.54 |  |  |  |
| caprylic (octanoic) acid | 1.53 |  | 1.22 | 1.24 |
| creatinine |  |  | 1.42 |  |
| cysteine |  |  | 1.57 | 0.61 |
| dodecanol | 2.03 |  |  | 1.62 |
| ethanolamine | 0.42 |  |  | 0.48 |
| fructose-6-phosphate | 2.13 |  | 2.23 |  |
| fucose | 1.61 |  | 4.07 |  |
| fumaric acid | 0.74 |  |  | 0.74 |
| glucose | 4.37 |  |  |  |
| glucose-6-phosphate | 2.56 |  | 2.50 |  |
| glutamine | 1.83 |  |  | 0.51 |
| glycerol-alpha-phosphate | 0.37 | 0.60 | 0.46 |  |
| glycolic acid |  |  |  | 1.43 |
| hexose-6-phosphate | 2.30 |  | 2.58 |  |
| hydroxylamine | 2.01 |  |  | 2.64 |
| hypoxanthine |  |  |  | 0.55 |
| inosine | 0.75 |  |  |  |
| isoleucine |  | 1.37 |  |  |
| lactic acid | 1.79 | 1.54 | 1.51 |  |
| lauric acid |  |  |  | 2.31 |
| leucine |  | 1.39 |  |  |
| lignoceric acid |  |  |  | 2.01 |
| linoleic acid | 0.14 | 0.33 | 0.20 | 0.30 |
| lysine |  |  | 0.60 |  |
| malic acid | 0.80 |  |  | 0.68 |
| mannitol | 1.74 |  |  | 2.01 |
| methionine |  |  |  | 0.77 |
| octadecanol | 2.43 |  |  | 1.95 |
| oleic acid | 0.18 | 0.36 | 0.18 |  |
| ornithine |  |  |  | 0.62 |
| oxalic acid | 1.69 |  | 1.26 |  |
| palmitic acid | 1.51 |  |  |  |
| palmitoleic acid |  | 0.60 |  |  |
| pelargonic acid | 1.56 |  |  | 1.72 |
| phosphate | 1.29 |  |  | 1.24 |
| pyruvic acid | 6.70 |  |  |  |
| ribulose-5-phosphate |  | 1.52 | 1.88 |  |
| serine |  | 1.52 | 1.56 |  |
| squalene | 0.66 | 0.69 | 0.72 |  |
| succinic acid | 1.84 |  |  |  |
| sucrose | 4.89 |  |  |  |
| taurine |  |  | 7.34 |  |
| threonine | 1.24 |  | 1.32 |  |
| tocopherol alpha- |  |  |  | 1.67 |
| tyrosine |  |  |  | 0.78 |
| valine |  | 1.30 |  |  |
| uracil |  |  |  | 0.70 |
| urea |  |  | 0.62 |  |
| uric acid | 1.76 |  |  | 0.70 |
| uridine | 1.73 |  |  | 2.15 |
| xanthine |  |  |  | 0.49 |
| xylitol |  |  | 0.49 | 0.62 |
| Number filled: | 37.00 | 14.00 | 26.00 | 32.00 |

### Tissue changes over the course of an infection

As described above, we collected tissue from mice infected for varying lengths of time. Initially, we hoped to compare all these samples, but were not able to do this because the metabolomics analyses were run on separate days and no normalization methods could remove the analysis-day bias. We therefore compared age-matched uninfected with 8 month-infected samples, and 3 day with 28 day infected samples because these sample metabolomes were analyzed together. Overall, the antral tissue remained stable, with no significant metabolite changes, between 0-8 months and 3-28 days. The only differences detected were in the corpus, specifically comparing uninfected tissue to 8 month infected tissue. In this case, there were 37 identified compounds that differed significantly (Table 7) and 61 total (Supplemental Table 8). Compounds that substantially increased during this time included a nearly 5-fold increase in adenosine, several lipids including those related to phospholipids (arachidonic acid, its product arachidic acid, oleic acid, linoleic acid), the lipid 1-monoolein, and the cholesterol precursor squalene. Additionally, there were ∼1.5-2-fold increases in the ATP product inosine, and the inorganic compound hydroxylamine. Compounds that decreased included the purine derivative uric acid, several amino acids (glutamine, aspartic acid, ornithine, aminomalonate) and the sugar fructose-6-phosphate. These data suggest that the antrum metabolome remains remarkably stable in the face of an *H. pylori* infection. The corpus, in contrast, displays several changes between the uninfected and chronically infected mice. These include indicators of tissue damage at the later time points, as well as depletion of key nutrients such as glutamine and fructose-6-phosphate that support cell proliferation.

**Table 7.** Differences in the corpus tissue between 0 and 8 months

| Metabolite | Fold change | FDR adjusted p |
| --- | --- | --- |
|  | Uninfected/8 mos | value: |
| uric acid | 0.22 | 0.001 |
| glutamine | 0.32 | 0.007 |
| fructose-6-phosphate | 0.35 | 0.037 |
| xanthine | 0.36 | 0.001 |
| aspartic acid | 0.39 | 0.004 |
| aminomalonate | 0.44 | 0.001 |
| ornithine | 0.46 | 0.001 |
| cysteine | 0.50 | 0.010 |
| fucose | 0.53 | 0.010 |
| urea | 0.55 | 0.002 |
| lactic acid | 0.57 | 0.010 |
| oxoproline | 0.58 | 0.005 |
| glycine | 0.58 | 0.001 |
| serine | 0.60 | 0.015 |
| methionine | 0.63 | 0.003 |
| alanine | 0.63 | 0.010 |
| threonine | 0.65 | 0.001 |
| **1,5-anhydroglucitol** | 0.70 | 0.033 |
| **creatinine** | 0.70 | 0.005 |
| **putrescine** | 0.72 | 0.010 |
| **valine** | 0.72 | 0.028 |
| **hypoxanthine** | 0.80 | 0.024 |
| **isoleucine** | 0.84 | 0.011 |
| **palmitic acid** | 1.27 | 0.013 |
| **phosphate** | 1.30 | 0.011 |
| **caprylic acid** | 1.32 | 0.031 |
| **stearic acid** | 1.35 | 0.035 |
| **arachidic acid** | 1.44 | 0.015 |
| **cholesterol** | 1.64 | 0.001 |
| **inosine** | 1.68 | 0.001 |
| **hydroxylamine** | 1.96 | 0.030 |
| **squalene** | 2.32 | 0.001 |
| **arachidonic acid** | 2.70 | 0.024 |
| **oleic acid** | 3.26 | 0.025 |
| **1-monoolein** | 3.26 | 0.019 |
| **linoleic acid** | 3.61 | 0.021 |
| **adenosine** | 4.88 | 0.001 |

## Discussion

We report the metabolomic differences that exist between gastric corpus and antral organoid-secreted products and gastric tissues, and how these two metabolomes respond to *H. pylori* infection. The gastric organoid data is the most straightforward to interpret because we were able to analyze the extracellular molecules. We find that *H. pylori* infection leads to depletion of specific metabolites in both the corpus and antrum, and these lost metabolites do not differ much between these regions. Our results suggest the preferred carbon and energy sources of *H. pylori* are the carboxylic acids lactic acid, lactamide, fumaric acid, malic acid and citric acid as well as the amino acids glutamine, serine, N-acetylglutamate, aspartic acid, oxoproline/pyroglutamic acid, and tyrosine. The simplest interpretation is that these compounds are used by *H. pylori*, an idea that is supported by the decrease in compounds that are known to be used by *H. pylori* such as urea, glutamine, cholesterol, lactic acid, and fumaric acid, although it’s possible that they are actively depleted by the mammalian cells after *H. pylori* infection.

Surprisingly, little is known about the preferred nutrient and energy sources of *H. pylori* when confronted with a complex mixture. Several studies support that the microbe does not substantially rely on sugars such as glucose, and instead relies on amino acids and carboxylic acids (27–29). Previous work supports the catabolism of carboxylic acids that we also show are used here. *H. pylori* can catabolize fumaric acid (30), malic acid (24), and lactic acid (22). Catabolism of carboxylic acids requires first their uptake. *H. pylori* has a C4-dicarboxylate transport protein that takes up fumaric acid and malic acid but not the larger citric acid (31). Citric acid complexes with iron, and there are several known ferric and ferrous citrate uptake systems in that have been characterized for their iron uptake properties but also presumably bring citrate into the cell as well (32). Another interpretation is that citric acid is depleted from the supernatant because it is used as a source of iron. Lactic acid has a separate uptake system, the LctP lactate permease. *H. pylori* actually has two lactate permease genes, *lctP1* (HP0140) and *lctP2* (HP0141), which are encoded next to each other in the genome (22). Studies have shown these two proteins are needed for lactate uptake, but have not yet determined if they play distinct roles.

Another finding from this work is that amino acids are depleted upon *H. pylori* infection. Amino acids can act as carbon, nitrogen, and energy sources. Work using defined medium determined that there are several *H. pylori* essential amino acids (e.g. arginine, histidine, leucine, methionine, phenylalanine, and valine), with others showing cross-study variation (alanine, cysteine, proline, serine and isoleucine) (23) but—for the most part—these were not the ones that were depleted here. Instead, it seems most likely that our depleted amino acids are carbon/nitrogen or energy sources. This idea agrees well with findings that *H. pylori* can catabolize amino acids as sole carbon/energy sources for *H. pylori* growth (29). Specifically, *H. pylori* catabolizes a large number of amino acids with the most substantial being alanine, arginine, asparagine, aspartate, glutamate, glutamine, proline, and serine, with studies reporting variation in the use of alanine, glycine, and valine (24, 29, 33, 34). This list is consistent with the amino acids depleted in this work although we did not detect depletion of all of these, and additionally found depletion of tyrosine and the glutamate derivatives N-acetylglutamate and pyroglutamic acid/oxoproline. *H. pylori* has several transporters that can take up amino acids including GltS for glutamate and DcuA for aspartate (35). These same transporters also take up glutamine and asparagine after they are deamidated to glutamate and aspartate. Other known transporters include those for peptides, Dpp and Opp (36, 37). There remain several unknowns. For serine, there is a predicted but as-yet uncharacterized serine uptake transporter (*sdaC*/HP0133), there is no predicted tyrosine transporter, and it is not yet known whether GltS can take up glutamate derivatives like N-acetylglutamate or pyroglutamic acid.

The mouse gastric tissue analysis uncovered that the antral metabolome is surprisingly stable over the course of an 8 month infection while the corpus showed significant changes. Another study analyzed the mouse gastric metabolome over an infection time course, but did not separate the corpus and the antrum (13). These authors also found relatively minimal changes in the mouse gastric metabolome over time. Our data suggest the infected corpus tissue has hallmarks of damage including elevated phospholipid products and adenosine, as well as depletion of important nutrients including the amino acids glutamine, aspartic acid and the sugar fructose-6-phosphate. *H. pylori* has been shown to use chemotaxis to preferentially colonize sites of gastric injury, further suggesting that these changes may lead to accumulation of *H. pylori* in the corpus region (38). Taken together, the metabolomics supports the idea that the corpus is undergoing atrophy, a known *H. pylori*-associated condition (39), and indeed that released phospholipids may be early indicators of carcinogenesis (40). Adenosine has immunosuppressive properties, and also has been associated with cancer development (41, 42). Thus the data might suggest that the *H. pylori* mouse model develops signs of carcinogenesis, cell damage, immunosuppression that can be detected using metabolomics.

In summary, our studies suggest that *H. pylori* utilizes multiple carboxylic acids and amino acids in both the corpus and the antral organoids, with minimal difference in metabolite usage between the two epithelia. We also find that there is very little overlap between the metabolites detected in the gastric organoid and tissue systems, reflecting the challenges of comparing the two systems. Lastly, we report that the antral metabolome is surprisingly stable over an 8 month *H. pylori* infection period, while the corpus displays significant changes that may indicate an early carcinogenic process.

## Acknowledgements

The authors are grateful to Eitaro Aihara and Chip Montrose (Univ. of Cincinnati) for gastric organoid culture training and advice; Andrew Liu and Kyle Kaminski (UC Santa Cruz) for experimental assistance; Kevin Johnson and Shuai Hu (UC Santa Cruz) for comments on the manuscript. The work described here was supported by National Institutes of Health National Institute of Allergy and Infectious Disease grant RO1AI116946 (to K.M.O.), the National Institutes of Health National Institute of Diabetes and Digestive and Kidney Diseases grant R01DK083402 (to Y.Z.), the California Cancer Research Coordinating Committee grant CRC-15-380545 (to K.M.O.) and a postdoctoral fellowship from Leopoldina (to D.K.). The funders had no role in study design, data collection and interpretation, or the decision to submit the work for publication.

## Material and Methods

### *H. pylori* strains and growth conditions

*H. pylori* strain SS1 with the GFP-expressing plasmid pTM115 was used for all studies (9, 43, 44). *H. pylori* were grown at 37°C, 5% O2, and 10% CO2 (balance N2). For solid media, we used Columbia blood agar (BD Diagnostics, Fisher Scientific)) with 5% difibrinated horse blood (Hemostat Labs, Dixon, California), 50 µg/ml cycloheximide (VWR), 10 µg/ml vancomycin, 5 µg/ml cefsulodin, 2.5 Units/ml polymyxin B (all from Gold Biotechnology, St Louis, MO), and 0.2% (wt/vol) β-cyclodextrin (Spectrum Labs, Gardena, CA) (CHBA). For liquid media, we used Brucella Broth (BBL) media supplemented with 10% heat-inactivated fetal bovine serum (FBS, Life Technologies) (BB10). For metabolomics analysis, *H. pylori* was grown for 24 hours in BB10 broth, and then filtered to remove bacterial cells as described below.

### Organoid preparation

All organoids and tissues were prepared from female C57Bl/6N mice (Helicobacter-free; Charles River). The University of California, Santa Cruz (UCSC) Institutional Animal Care and Use Committee approved all animal protocols and experiments (Protocol #OTTEK1505). All animal procedures were in strict accordance with the NIH Guide for the Care and Use of Laboratory Animals. Mice were housed at the UCSC animal facility and given free access to food and water. For gastric organoids, 12 week old mice were sacrificed by CO2 narcosis, and the stomach was removed by cutting at the stomach-esophageal junction and the antrum-duodenum sphincter. The forestomach was removed and the stomach opened along the lesser curvature using scissors. The stomach was gently rinsed by moving back and forth in 25 ml ice cold phosphate buffered saline (PBS). Blood vessels and the muscle layer were removed from the non-luminal side of the tissue using microscissors (Kelly Scientific) and tweezers. The corpus was separated from the antrum with a scalpel, based on the difference in tissue coloration as a marker for the border between these regions, excluding a ∼ 5mm section between the two tissues to ensure purity. Each piece was further divided into thirds to be used for organoid preparation, bacterial enumeration, or flash frozen in liquid N2 for metabolomics analysis.

For organoids, we next isolated using a protocol adapted from Mahe *et al*. (45). Dissected gastric tissue was cut into 1 mm^2^ pieces and incubated with slight shaking in Dulbeccos Phosphate buffered saline (DPBS, Millipore) with 5mM ETDA at 4°C for 2 hours. After this period, the tissue was transferred into ice-cold DBPS containing 1% sucrose and 1.5% sorbitol, and shaken roughly by hand for 2 minutes. The remaining large tissue pieces were allowed to settle, and 2 ml of the solution containing the glands was removed. Glands were washed with cold DPBS, and mixed into 50 µl of matrigel (BD Biosciences). Matrigel and glands were placed into a 24-well plate if being prepared for flat organoids and incubated for 10 minutes at 37°C, until the Matrigel solidified. 3D-gastric organoid media was then added, to cover the Matrigel and the glands. 3D-gastric organoid media contains Advanced DMEM/F12 (Invitrogen) supplemented with 2mM GlutaMax (Invitrogen), 10 mM HEPES (Sigma), 100 U/mL penicillin/100 μg/mL streptomycin (Invitrogen), 1× N2 supplement (Invitrogen), 1× B27 supplement (Invitrogen), 1 μg/mL mouse R-spondin (R&D systems), 100 ng/mL Noggin (Peprotech), 50 ng/mL Epidermal Growth Factor (EGF, Peprotech). Media was exchanged with fresh media every 2 days for up to 7 days.

To create 2D organoids, the spheroid gastric organoids were collected with ice-cold DPBS, the matrigel removed by a washing and centrifugation at 195xg for 5 minutes at 4°C. Organoids were collected and resuspended in 500 µl 2D culture media (DMEMF12, Advanced-Base Medium, 10% Fetal Bovine Serum, 10 mM HEPES, 2mM Glutamine, 1X N2 supplement, 1X B27 supplement, 10 µM Y-compound (Y-27632, Sigma), 50 ng/ml EGF) and placed into a collagen-coated 24 well plate for attachment. After 24 hours, the media was replaced with fresh 2D culture media plus 1×10^7 *H. pylori* SS1 pTM115 per well. The *H. pylori* was grown in BB10 as above. After 24 hours of infection, media was collected from the organoids or bacterial cells, passed through a 0.2 µm filter to remove organoid and bacterial cells from the media, and quick frozen in liquid N2. For this analysis, we had 6 independent samples for most groups (corpus infected, corpus uninfected, antrum infected, antrum uninfected and the *H. pylori* alone control) and 3 independent samples for the media alone.

### Mouse tissue collection and preparation

Female C57Bl6/N mice (Charles River) were infected at 4-6 weeks of age with 500 µl *H. pylori* SS1 pTM115 grown in BB10 media to mid-exponential phase (OD600 ∼0.3). Infection was accomplished by oral infection with a 20-gauge × 1.5 inch feeding needle (Popper). At the desired time, the mouse stomachs were collected as for organoids and divided into corpus and antrum. For metabolomics, pieces were flash frozen in liquid N2. For bacterial number determination, the piece was weighed, homogenized using the Bullet Blender (Next Advance) with 1.0-mm zirconium silicate beads, diluted, and plated on CHBA with the addition of 20 µg/ml bacitracin, 10 µg/ml nalidixic acid, and 15 µg/ml kanamycin.

### Metabolomic profiling

Samples for metabolomics were quick frozen in liquid N2, stored at -80C, and shipped on dry ice to the UC Davis NIH West Coast Metabolomics Center (https://metabolomics.ucdavis.edu). An untargeted analysis of primary metabolites was performed using automatic liner exchange chromatography and cold injection gas chromatography (GC) time of flight (TOF) mass spectrometry methods as detailed in Fiehn *et al.* (46). Briefly, the GC column was a GC Restek corporation Rtx-5Sil MS column with a mobile phase of helium with automatic liner exchanges after each set of 10 injections. Mass spectrometry parameters were as follows: a Leco Pegasus IV mass spectrometer is used with unit mass resolution at 17 spectra s-1 from 80-500 Da at -70 eV ionization energy and 1800 V detector voltage with a 230°C transfer line and a 250°C ion source. ChromaTOF vs. 2.32 was used for data preprocessing without smoothing, 3 s peak width, baseline subtraction just above the noise level, and automatic mass spectral deconvolution and peak detection at signal/noise levels of 5:1 throughout the chromatogram. Apex masses are reported for use in the BinBase algorithm. Result *.txt files are exported to a data server with absolute spectra intensities and further processed by a filtering algorithm implemented in the metabolomics BinBase database. The BinBase algorithm (rtx5) used the settings: validity of chromatogram (<10 peaks with intensity >10^7 counts s-1), unbiased retention index marker detection (MS similarity>800, validity of intensity range for high m/z marker ions), retention index calculation by 5th order polynomial regression. Spectra are cut to 5% base peak abundance and matched to database entries from most to least abundant spectra using the following matching filters: retention index window ±2,000 units (equivalent to about ±2 s retention time), validation of unique ions and apex masses (unique ion must be included in apexing masses and present at >3% of base peak abundance), mass spectrum similarity must fit criteria dependent on peak purity and signal/noise ratios and a final isomer filter. Failed spectra are automatically entered as new database entries if s/n >25, purity <1s.0 and presence in the biological study design class was >80%. All thresholds reflect settings for ChromaTOF v. 2.32. Quantitative data are reported as relative peak intensities (for example, normalized to the peak intensities found in quality control samples). Data is given with the following information: (1) Binbase name, which includes a name if the peak has been identified, or just a Binbase number/BBID if it is not identified; (2) Retention index (ret index); (3) ‘quant mz’ column details the m/z value that was used to quantify the peak height of each entry.(4) The ‘mass spec’ column details the complete mass spectrum of the metabolite given as mz: intensity values, separated by spaces. (5) for identified compounds, external database identifiers such as InChI key, defined by the IUPAC and NIST consortia; PubChem ID, the unique identifier of a metabolite in the PubChem database; and KEGG ID gives the unique identifier associated with an identified metabolite in the community database KEGG LIGAND DB. The ‘internal standard’ addition within the BinBase name clarifies if a specific chemical has been added into the extraction solvent as internal standard. These internal standards serve as retention time alignment markers, for quality control purposes and for quantification corrections. The full organoid metabolomics data is given in Supplemental Data Set 1 and the full mouse tissue metabolomics is given as Supplemental Data Set 2.

### Metabolomics Statistical Analysis

For normalization, a variant of a ‘vector normalization’ in which the sum of all peak heights for all <u>identified</u> metabolites was calculated for each sample to create the value called mTIC (47). All mTIC averages were then compared. If these averages differ by p<0.05, data was normalized to the average mTIC of each group; this was the case for the mouse tissue data. If averages between treatment groups are not different, data was normalized to the total average mTIC; this was the case for the organoid data. For principal component analysis (PCA), we applied log10 transformation and auto-scaling to the data. Difference comparisons between multiple groups was made using a one-way analysis of variance (ANOVA) or between two groups using the Students T test. To control the false discovery rate (FDR), we adjusted the p-values using Benjamini-Hochberg procedure (48). The mean fold change was calculated for each pair of two groups using non-transformed data. To generate the cluster analysis heat map shown in Fig. 2, the known chemicals peak height from the organoid data was submitted to Metaboanalyst (https://www.metaboanalyst.ca/) heatmap and clustering analysis. The following parameters were used: concentrations, samples in columns unpaired, data normalization by sum, and auto-scaling. For output, we chose the top 40 by ANOVA.

### Enrichment Analysis

Chemical similarly enrichment analysis was performed using the ChemRICH approach (http://chemrich.us) (17)with the Mann-Whitney U test p-values and median fold-changes of our identified metabolites. ChemRICH was implemented using the Kolmogorov–Smirnov test on the identified clusters to evaluate whether a metabolite cluster was represented more than expected by chance.

---

**Supplemental Fig. 1.**
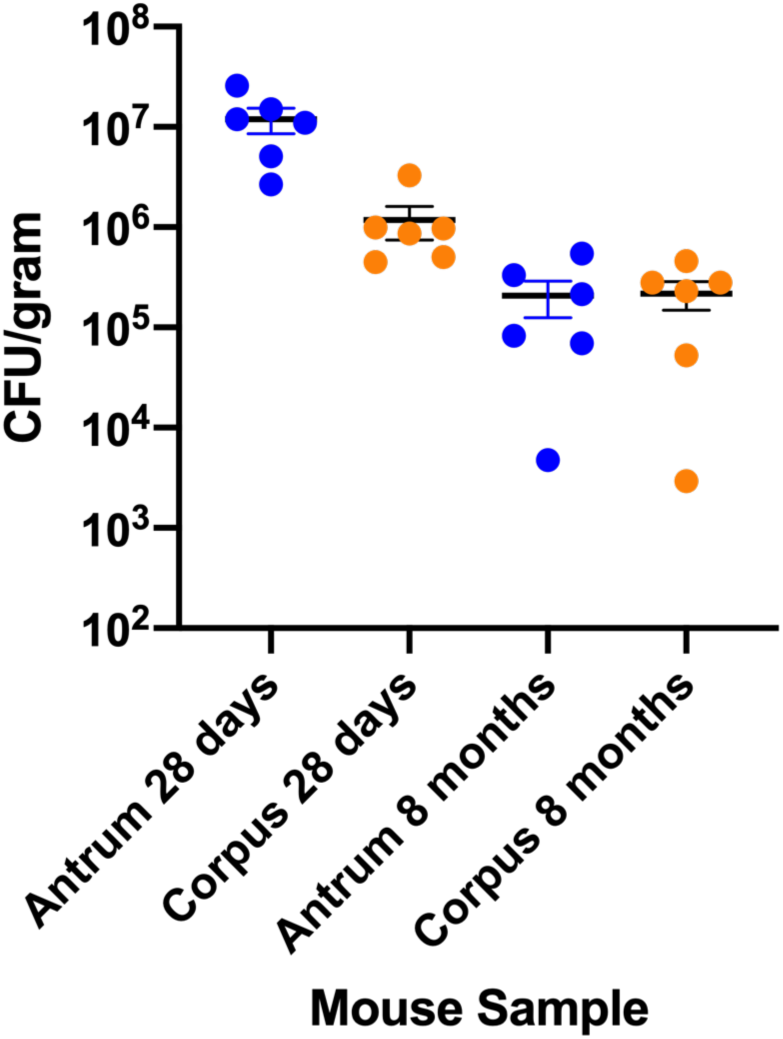
*H. pylori* SS1 pTM115 infection levels were determined by homogenizing and plating *H. pylori* tissue samples on CHBA plates, as described in (9).

**Supplemental Table 1.** All compounds detected in corpus and antrum organoids, ordered by most significantly different in the uninfected tissue.

| Metabolite | Uninfected Antrum * Corpus<br>(Adjusted P) | Infected Antrum *<br>Corpus (adjusted P) |
| --- | --- | --- |
| phosphoethanolamine | 0.002651985 | 0.000180099 |
| uridine | 0.005419144 | 0.464113525 |
| lactic acid | 0.007430126 | 0.000180099 |
| 104906 | 0.017715615 | 0.086153073 |
| malic acid | 0.017715615 | 0.268980032 |
| glycerol | 0.017715615 | 0.582983261 |
| alanine | 0.02691136 | 0.205852591 |
| isoleucine | 0.040300273 | 0.836884308 |
| 1029 | 0.047459149 | 0.231705706 |
| fumaric acid | 0.07430243 | 0.356836096 |
| aspartic acid | 0.157896458 | 0.086153073 |
| 5-methoxytryptamine | 0.205835259 | 0.284930795 |
| leucine | 0.205835259 | 0.81471968 |
| serine | 0.210671869 | 0.284930795 |
| 110628 | 0.223964174 | 0.191110038 |
| 31272 | 0.238827659 | 0.191110038 |
| 110624 | 0.238827659 | 0.231705706 |
| beta-sitosterol | 0.238827659 | 0.242916418 |
| 21704 | 0.238827659 | 0.343995331 |
| 2011 | 0.238827659 | 0.505913205 |
| succinic acid | 0.238827659 | 0.517839725 |
| 171297 | 0.238827659 | 0.728199809 |
| valine | 0.238827659 | 0.782301967 |
| 124844 | 0.238827659 | 0.936919893 |
| tyrosine | 0.254154621 | 0.51753482 |
| citric acid | 0.258763862 | 0.636757531 |
| 2-ketoisocaproic acid | 0.276555644 | 0.642081065 |
| 84213 | 0.293241669 | 0.018516393 |
| 18488 | 0.293241669 | 0.22578453 |
| 592 | 0.293241669 | 0.284260336 |
| 4937 | 0.293241669 | 0.446579436 |
| 1-monoolein | 0.293241669 | 0.482490002 |
| **1702** | 0.293241669 | 0.515564206 |
| **106936** | 0.293241669 | 0.519845974 |
| **ethanolamine** | 0.293241669 | 0.782301967 |
| **146957** | 0.293241669 | 0.81471968 |
| **453** | 0.293241669 | 0.81471968 |
| **1721** | 0.293241669 | 0.946540754 |
| **4901** | 0.293241669 | 0.968800142 |
| **110607** | 0.31434439 | 0.191110038 |
| **glutamine** | 0.328468904 | 0.908297593 |
| **hypoxanthine** | 0.332745395 | 0.065661824 |
| **pyruvic acid** | 0.332745395 | 0.071370692 |
| **110611** | 0.332745395 | 0.284930795 |
| **palmitoleic acid** | 0.332745395 | 0.512257418 |
| **41891** | 0.332745395 | 0.936919893 |
| **citramalic acid** | 0.335550733 | 0.007095858 |
| **2438** | 0.337553399 | 0.191110038 |
| **43100** | 0.337553399 | 0.22578453 |
| **threonine** | 0.337553399 | 0.549194862 |
| **4746** | 0.356526648 | 0.812487739 |
| **34001** | 0.377827315 | 0.631327492 |
| **134408** | 0.380200283 | 0.510040491 |
| **phenylalanine** | 0.380200283 | 0.572210196 |
| **glutamate** | 0.384886914 | 0.836884308 |
| **84181** | 0.400166069 | 0.643917535 |
| **117998** | 0.406236611 | 0.618230449 |
| **palmitic acid** | 0.423562874 | 0.715981087 |
| **6767** | 0.426301631 | 0.936919893 |
| **xylose** | 0.439560043 | 0.83201964 |
| **stearic acid** | 0.443205758 | 0.836884308 |
| **glucose** | 0.459811251 | 0.231705706 |
| **84088** | 0.459811251 | 0.284930795 |
| **95422** | 0.459811251 | 0.333148932 |
| **guanosine** | 0.478340166 | 0.284260336 |
| **glycine** | 0.478340166 | 0.436484516 |
| **1684** | 0.478340166 | 0.836110331 |
| **linoleic acid** | 0.478340166 | 0.87510423 |
| **110610** | 0.480381473 | 0.231705706 |
| **110604** | 0.480381473 | 0.923206508 |
| **307** | 0.484483702 | 0.367975944 |
| 110268 | 0.484483702 | 0.936919893 |
| 106629 | 0.507178508 | 0.691984508 |
| 1,5-anhydroglucitol | 0.510210459 | 0.975336286 |
| 39 | 0.518000515 | 0.59526644 |
| 46128 | 0.519371643 | 0.924926144 |
| 42525 | 0.529503528 | 0.631327492 |
| 4793 | 0.537723431 | 0.936919893 |
| 110606 | 0.537723431 | 0.970472622 |
| heptadecanoic acid | 0.553906672 | 0.833861163 |
| 104179 | 0.563207313 | 0.280690651 |
| 21706 | 0.563207313 | 0.296516958 |
| oxalic acid | 0.563207313 | 0.790834054 |
| 121002 | 0.563207313 | 0.81471968 |
| 462 | 0.563207313 | 0.857100411 |
| lignoceric acid | 0.563207313 | 0.989881797 |
| n-acetylglutamate | 0.565556245 | 0.313838668 |
| UDP-glucuronic acid | 0.565556245 | 0.482490002 |
| 137 | 0.565556245 | 0.515564206 |
| 18305 | 0.565556245 | 0.587183915 |
| 121890 | 0.565556245 | 0.686669057 |
| 257 | 0.565556245 | 0.774723994 |
| 62304 | 0.565556245 | 0.836884308 |
| 5242 | 0.566168823 | 0.933465375 |
| 171305 | 0.579019438 | 0.328437176 |
| 129225 | 0.579019438 | 0.918317387 |
| oxoproline | 0.579019438 | 0.985655378 |
| 104131 | 0.592096735 | 0.642081065 |
| 91421 | 0.592096735 | 0.654916617 |
| 16747 | 0.595132533 | 0.191110038 |
| 1-monostearin | 0.595132533 | 0.505913205 |
| 126397 | 0.595132533 | 0.734162534 |
| 3361 | 0.595132533 | 0.736200584 |
| asparagine | 0.595132533 | 0.789090761 |
| 42913 | 0.595132533 | 0.864098289 |
| ribulose-5-phosphate | 0.595132533 | 0.896738468 |
| inosine | 0.595132533 | 0.927157981 |
| 133514 | 0.595132533 | 0.970472622 |
| 4766 | 0.601231218 | 0.81471968 |
| uracil | 0.603297379 | 0.007095858 |
| **110315** | 0.603297379 | 0.512257418 |
| **ribose** | 0.603297379 | 0.691984508 |
| **5085** | 0.603297379 | 0.970472622 |
| **88583** | 0.610618327 | 0.766586764 |
| **7490** | 0.614657762 | 0.284930795 |
| **84383** | 0.631546366 | 0.877738085 |
| **5837** | 0.634587824 | 0.284930795 |
| **816** | 0.634587824 | 0.936919893 |
| **109981** | 0.634587824 | 0.950234967 |
| **3-hydroxybutyric acid** | 0.646217257 | 0.231705706 |
| **98** | 0.646217257 | 0.338345742 |
| **124826** | 0.646217257 | 0.686730698 |
| **39801** | 0.646217257 | 0.857100411 |
| **4976** | 0.651752253 | 0.715981087 |
| **nicotinamide** | 0.656059674 | 0.367975944 |
| **sucrose** | 0.675162143 | 0.462239268 |
| **sorbitol** | 0.675162143 | 0.808176059 |
| **47358** | 0.675162143 | 0.81471968 |
| **myo-inositol** | 0.675162143 | 0.827285838 |
| **105554** | 0.675162143 | 0.936919893 |
| **50367** | 0.679604667 | 0.78171695 |
| **97455** | 0.679604667 | 0.946540754 |
| **17651** | 0.701216227 | 0.018073951 |
| **299** | 0.701216227 | 0.22578453 |
| **capric acid** | 0.701657334 | 0.231705706 |
| **squalene** | 0.701657334 | 0.512257418 |
| **urea** | 0.750285799 | 0.284930795 |
| **133242** | 0.750285799 | 0.287308954 |
| **110618** | 0.751982904 | 0.284260336 |
| **103886** | 0.751982904 | 0.519845974 |
| **110284** | 0.751982904 | 0.579843648 |
| **arachidonic acid** | 0.751982904 | 0.737061058 |
| **threonic acid** | 0.751982904 | 0.78171695 |
| **171823** | 0.751982904 | 0.81471968 |
| **pantothenic acid** | 0.751982904 | 0.836110331 |
| **ornithine** | 0.751982904 | 0.908297593 |
| **adenosine** | 0.751982904 | 0.923068234 |
| **128034** | 0.753901495 | 0.086153073 |
| **2242** | 0.753901495 | 0.81471968 |
| **1704** | 0.753901495 | 0.895894859 |
| **methionine** | 0.756140424 | 0.314027382 |
| **124903** | 0.756140424 | 0.435620298 |
| **131620** | 0.756140424 | 0.505913205 |
| **glucose-6-phosphate** | 0.756140424 | 0.506557439 |
| **pelargonic acid** | 0.756140424 | 0.734162534 |
| **84116** | 0.756140424 | 0.963604634 |
| **18520** | 0.762520561 | 0.231705706 |
| **lyxitol** | 0.763146274 | 0.528294835 |
| **benzoic acid** | 0.763442027 | 0.242916418 |
| **1872** | 0.769085302 | 0.284930795 |
| **106693** | 0.769085302 | 0.416839778 |
| **beta-alanine** | 0.769085302 | 0.927157981 |
| **124825** | 0.769085302 | 0.946540754 |
| **110550** | 0.769085302 | 0.968800142 |
| **31563** | 0.779838994 | 0.22578453 |
| **2061** | 0.779838994 | 0.242916418 |
| **133244** | 0.779838994 | 0.284930795 |
| **47** | 0.779838994 | 0.333148932 |
| **84682** | 0.779838994 | 0.605733645 |
| **4797** | 0.779838994 | 0.631327492 |
| **102716** | 0.779838994 | 0.886543646 |
| **isothreonic acid** | 0.785478385 | 0.946540754 |
| **1708** | 0.786113038 | 0.462239268 |
| **fucose** | 0.786113038 | 0.578472168 |
| **5269** | 0.786113038 | 0.636757531 |
| **111776** | 0.786113038 | 0.720121509 |
| **cystine** | 0.786133924 | 0.288992597 |
| **octadecanol** | 0.786133924 | 0.45964984 |
| **levoglucosan** | 0.786133924 | 0.482490002 |
| **pseudo uridine** | 0.786133924 | 0.578472168 |
| **121028** | 0.786133924 | 0.636757531 |
| **trans-4-hydroxyproline** | 0.786133924 | 0.836884308 |
| **N-acetylmannosamine** | 0.786133924 | 0.836884308 |
| **133348** | 0.786133924 | 0.946540754 |
| **1713** | 0.792617072 | 0.789090761 |
| **108936** | 0.811127089 | 0.284260336 |
| **hexitol** | 0.811127089 | 0.284930795 |
| **171564** | 0.811127089 | 0.512257418 |
| **146430** | 0.811127089 | 0.598662621 |
| **172625** | 0.811127089 | 0.946540754 |
| **methanolphosphate** | 0.811127089 | 0.985655378 |
| **taurine** | 0.81569458 | 0.38642342 |
| **hexose-6-phosphate** | 0.818819529 | 0.231705706 |
| **124881** | 0.825851935 | 0.312885721 |
| **adenosine-5-monophosphate** | 0.825851935 | 0.482490002 |
| **100253** | 0.825851935 | 0.517839725 |
| **270** | 0.825851935 | 0.568298427 |
| **1709** | 0.825851935 | 0.628058033 |
| **112533** | 0.825851935 | 0.812487739 |
| **171303** | 0.825851935 | 0.836110331 |
| **fructose** | 0.825851935 | 0.836884308 |
| **4945** | 0.83000812 | 0.782301967 |
| **102711** | 0.83000812 | 0.968800142 |
| **17068** | 0.83715613 | 0.528294835 |
| **creatinine** | 0.839148162 | 0.836884308 |
| **31359** | 0.839285803 | 0.284260336 |
| **34089** | 0.839285803 | 0.691567949 |
| **171970** | 0.839285803 | 0.774723994 |
| **62** | 0.839285803 | 0.946540754 |
| **110** | 0.839334928 | 0.517839725 |
| **glyceric acid** | 0.839334928 | 0.823780019 |
| **97332** | 0.839334928 | 0.886543646 |
| **myristic acid** | 0.85588809 | 0.578472168 |
| **99** | 0.85617772 | 0.636757531 |
| **48428** | 0.87391282 | 0.836884308 |
| **lactamide** | 0.87416202 | 0.005989406 |
| **glycerol-alpha-phosphate** | 0.87416202 | 0.231705706 |
| **42007** | 0.87416202 | 0.284930795 |
| **31357** | 0.87416202 | 0.365962254 |
| **110321** | 0.87416202 | 0.536608647 |
| **2575** | 0.87416202 | 0.636757531 |
| **135610** | 0.87416202 | 0.737061058 |
| **124868** | 0.87416202 | 0.812487739 |
| **47262** | 0.87416202 | 0.836884308 |
| **N-methylalanine** | 0.87416202 | 0.836884308 |
| **135384** | 0.87416202 | 0.933465375 |
| **26062** | 0.87416202 | 0.936919893 |
| **490** | 0.87416202 | 0.946540754 |
| **1-monopalmitin** | 0.875212416 | 0.818181154 |
| **glycolic acid** | 0.875212416 | 0.832383276 |
| **aminomalonate** | 0.875212416 | 0.936919893 |
| **1173** | 0.889444819 | 0.393398981 |
| **17253** | 0.889788777 | 0.284930795 |
| **119025** | 0.889788777 | 0.568298427 |
| **892** | 0.889788777 | 0.636757531 |
| **thymine** | 0.889788777 | 0.809589193 |
| **1981** | 0.889788777 | 0.946540754 |
| **caprylic acid** | 0.898189959 | 0.342276844 |
| **tocopherol alpha-** | 0.898189959 | 0.515564206 |
| **42914** | 0.898189959 | 0.51753482 |
| **127640** | 0.898189959 | 0.683562764 |
| **1912** | 0.898189959 | 0.700163732 |
| **172624** | 0.90612736 | 0.268980032 |
| **135260** | 0.916141921 | 0.968800142 |
| **41989** | 0.918622025 | 0.505913205 |
| **54** | 0.928348555 | 0.367975944 |
| **122191** | 0.928348555 | 0.450712839 |
| **42003** | 0.928348555 | 0.515564206 |
| **161878** | 0.929313 | 0.517839725 |
| **4546** | 0.929313 | 0.678307385 |
| **88502** | 0.930909724 | 0.231705706 |
| **3781** | 0.930909724 | 0.578472168 |
| **ribitol** | 0.930909724 | 0.582983261 |
| **tryptophan** | 0.930909724 | 0.695679906 |
| **121402** | 0.930909724 | 0.860865955 |
| **glucose-1-phosphate** | 0.932457697 | 0.642081065 |
| **84922** | 0.932457697 | 0.756172104 |
| **lauric acid** | 0.932457697 | 0.81471968 |
| **erythritol** | 0.932457697 | 0.836884308 |
| **21683** | 0.932457697 | 0.836884308 |
| **124870** | 0.932457697 | 0.936919893 |
| **maltose** | 0.941148991 | 0.981645483 |
| **1941** | 0.947643987 | 0.519845974 |
| **18252** | 0.947643987 | 0.836884308 |
| **46292** | 0.947775612 | 0.231705706 |
| arachidic acid | 0.947775612 | 0.923206508 |
| 108950 | 0.947775612 | 0.946540754 |
| 47170 | 0.95069283 | 0.81471968 |
| 1862 | 0.951213059 | 0.284930795 |
| hydroxylamine | 0.951213059 | 0.51753482 |
| 17664 | 0.951213059 | 0.631327492 |
| inositol-4-monophosphate | 0.951213059 | 0.642081065 |
| 170841 | 0.951213059 | 0.913905374 |
| 5990 | 0.952898391 | 0.634108425 |
| xylitol | 0.952898391 | 0.81471968 |
| 4531 | 0.95541352 | 0.946540754 |
| 6066 | 0.957595176 | 0.268980032 |
| 171285 | 0.957595176 | 0.496561445 |
| galactinol | 0.957595176 | 0.886543646 |
| oleic acid | 0.957595176 | 0.902484377 |
| 17044 | 0.957595176 | 0.936919893 |
| gluconic acid | 0.96053486 | 0.242916418 |
| 1878 | 0.96053486 | 0.284930795 |
| 103102 | 0.96053486 | 0.482490002 |
| 41811 | 0.96053486 | 0.756172104 |
| 132143 | 0.96053486 | 0.837533359 |
| 16572 | 0.96053486 | 0.946540754 |
| 1700 | 0.96053486 | 0.971569691 |
| 110355 | 0.965659677 | 0.915047353 |
| mannitol | 0.971697205 | 0.578472168 |
| methionine sulfoxide | 0.971697205 | 0.601233104 |
| 100730 | 0.981967674 | 0.231705706 |
| cholesterol | 0.981967674 | 0.280690651 |
| 34059 | 0.981967674 | 0.284260336 |
| homoserine | 0.981967674 | 0.284930795 |
| uric acid | 0.981967674 | 0.296516958 |
| 22423 | 0.981967674 | 0.367975944 |
| fructose-6-phosphate | 0.981967674 | 0.367975944 |
| threitol | 0.981967674 | 0.38642342 |
| 657 | 0.981967674 | 0.397180154 |
| 5576 | 0.981967674 | 0.438967876 |
| UDP-N-acetylglucosamine | 0.981967674 | 0.446579436 |
| 3-phosphoglycerate | 0.981967674 | 0.446579436 |
| thymidine | 0.981967674 | 0.462239268 |
| 1735 | 0.981967674 | 0.505913205 |
| 120744 | 0.981967674 | 0.517839725 |
| 31285 | 0.981967674 | 0.530137261 |
| 41808 | 0.981967674 | 0.555578447 |
| 6104 | 0.981967674 | 0.578472168 |
| 134336 | 0.981967674 | 0.642081065 |
| 10962 | 0.981967674 | 0.654916617 |
| 111263 | 0.981967674 | 0.664222466 |
| lysine | 0.981967674 | 0.691984508 |
| 5346 | 0.981967674 | 0.691984508 |
| 573 | 0.981967674 | 0.691984508 |
| 2,5-dihydroxypyrazine<br>NIST | 0.981967674 | 0.717523417 |
| histidine | 0.981967674 | 0.72318169 |
| 3258 | 0.981967674 | 0.734162534 |
| trehalose | 0.981967674 | 0.737061058 |
| 134 | 0.981967674 | 0.774723994 |
| hydroxycarbamate NIST | 0.981967674 | 0.790834054 |
| 2-monopalmitin | 0.981967674 | 0.790834054 |
| phosphate | 0.981967674 | 0.81471968 |
| 1875 | 0.981967674 | 0.81471968 |
| 130396 | 0.981967674 | 0.833861163 |
| dodecanol | 0.981967674 | 0.836884308 |
| beta-glycerolphosphate | 0.981967674 | 0.836884308 |
| 41683 | 0.981967674 | 0.844644327 |
| n-acetyl-d-hexosamine | 0.981967674 | 0.84921018 |
| xanthine | 0.981967674 | 0.864098289 |
| 1746 | 0.981967674 | 0.880867478 |
| 42424 | 0.981967674 | 0.91312373 |
| 120562 | 0.981967674 | 0.918317387 |
| 1064 | 0.981967674 | 0.927157981 |
| pyrophosphate | 0.981967674 | 0.944180589 |
| 133907 | 0.981967674 | 0.946540754 |
| 121404 | 0.981967674 | 0.963604634 |
| 443 | 0.981967674 | 0.989881797 |
| 106498 | 0.983353659 | 0.435620298 |
| 2-monoolein | 0.983353659 | 0.981645483 |
| 23635 | 0.983699575 | 0.284930795 |
| cysteine | 0.988054349 | 0.249768926 |
| **21666** | 0.988054349 | 0.936919893 |
| **133712** | 0.988054349 | 0.946540754 |
| **putrescine** | 0.989151077 | 0.880867478 |
| **103870** | 0.993269431 | 0.284260336 |
| **2-hydroxyglutaric acid** | 0.993882527 | 0.191110038 |
| **2936** | 0.993882527 | 0.284260336 |
| **170728** | 0.993882527 | 0.496411576 |
| **1717** | 0.993882527 | 0.636757531 |
| **41985** | 0.993882527 | 0.75375956 |
| **31224** | 0.993882527 | 0.902484377 |

**Supplemental Table 2.**
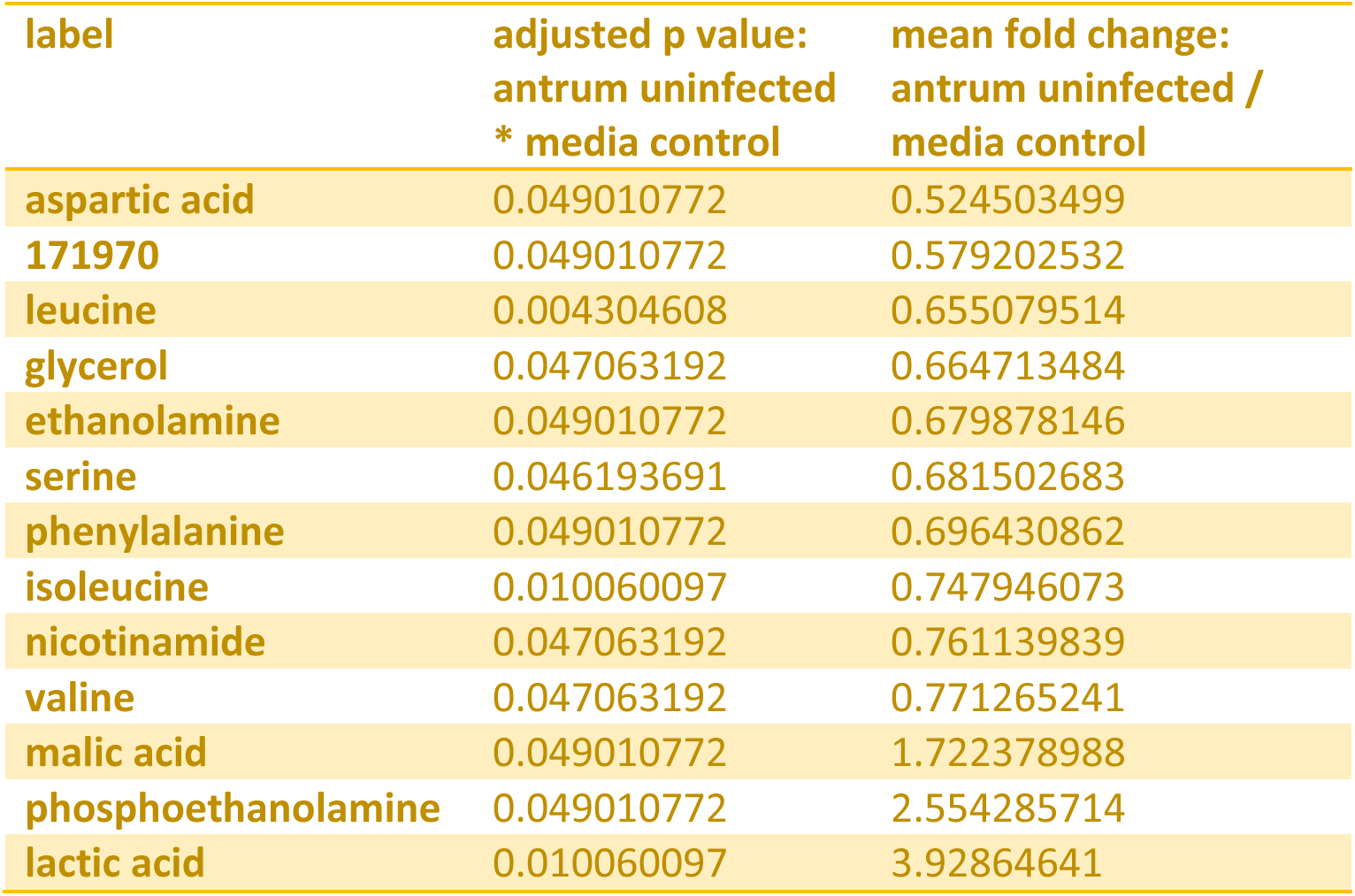
Compounds that differed between antrum organoids and media alone

**Supplemental Table 3.**
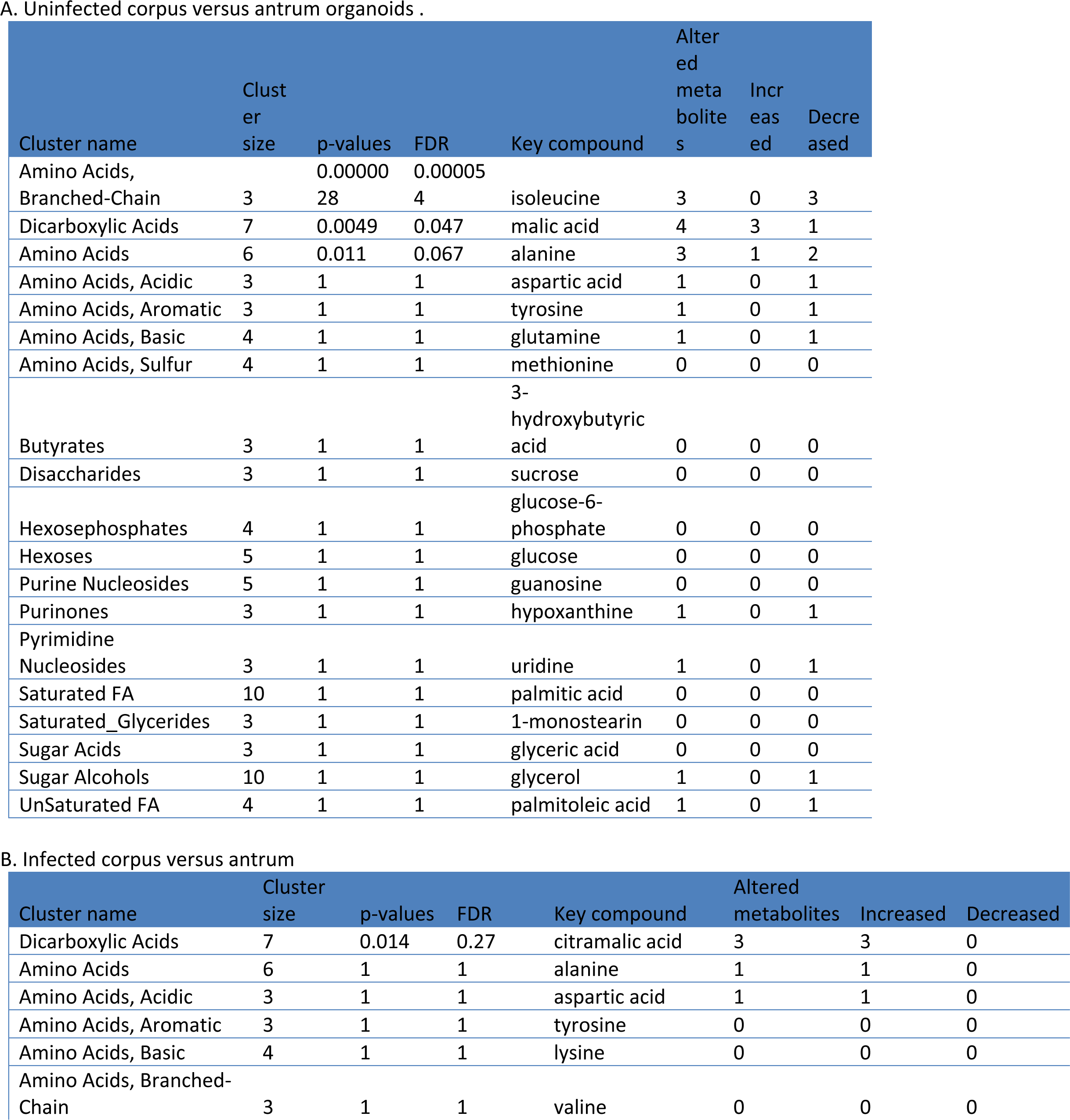
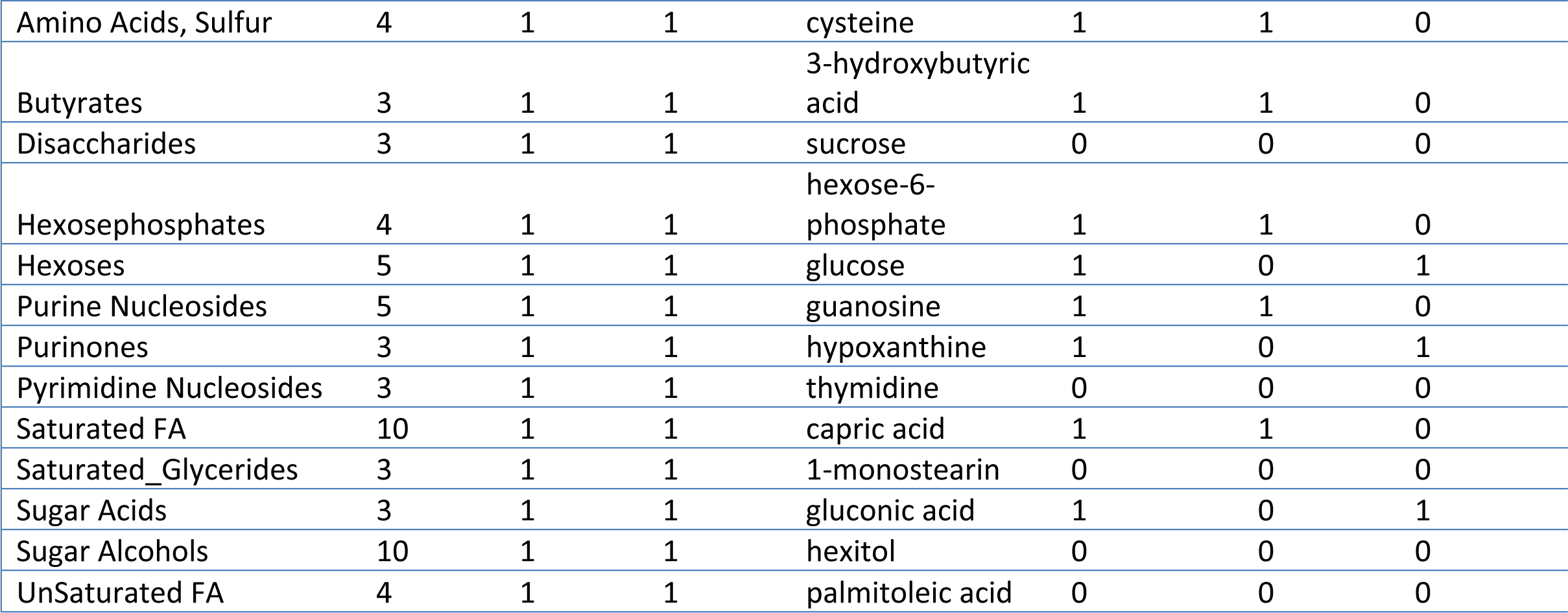
Enrichment analysis comparing corpus and antrum in uninfected and infected tissues.

**Supplemental Table 4.** All compounds that change in the antrum organoids upon infection, ordered by P value

| Label | adjusted p value: antrum<br>infected * antrum<br>uninfected |
| --- | --- |
| 21706 | 2.70849E-08 |
| 1708 | 2.70849E-08 |
| 1746 | 2.96284E-08 |
| urea | 5.7285E-08 |
| glutamate | 1.26084E-07 |
| 128034 | 1.26084E-07 |
| citric acid | 3.77645E-07 |
| 1721 | 4.8323E-07 |
| 1713 | 8.86461E-07 |
| 18520 | 1.11747E-06 |
| uridine | 3.06326E-06 |
| 133907 | 3.06326E-06 |
| 1717 | 4.15844E-06 |
| 270 | 4.92879E-06 |
| malic acid | 9.29347E-06 |
| ribose | 1.20005E-05 |
| leucine | 1.22013E-05 |
| 109981 | 1.22013E-05 |
| homoserine | 1.3636E-05 |
| 34001 | 1.3636E-05 |
| 4746 | 1.3636E-05 |
| succinic acid | 1.37595E-05 |
| 108950 | 1.69307E-05 |
| alanine | 1.69441E-05 |
| glycerol | 2.00477E-05 |
| 135260 | 2.65798E-05 |
| 84213 | 3.49234E-05 |
| cystine | 5.53993E-05 |
| uric acid | 6.01137E-05 |
| thymidine | 6.01137E-05 |
| 170728 | 6.01137E-05 |
| serine | 6.78232E-05 |
| histidine | 7.23439E-05 |
| phenylalanine | 8.76571E-05 |
| fumaric acid | 9.75406E-05 |
| glutamine | 9.75978E-05 |
| citramalic acid | 9.75978E-05 |
| 31272 | 9.79653E-05 |
| pyruvic acid | 0.000101866 |
| asparagine | 0.000110508 |
| 110355 | 0.000150299 |
| 97332 | 0.000193405 |
| 110315 | 0.000336396 |
| 1700 | 0.000351946 |
| n-acetylglutamate | 0.000363761 |
| threonine | 0.000397385 |
| glycine | 0.000636786 |
| 453 | 0.0006819 |
| 34059 | 0.000790494 |
| valine | 0.000912187 |
| 84088 | 0.001025989 |
| 43100 | 0.001142457 |
| 3361 | 0.001142457 |
| lysine | 0.001201798 |
| 171970 | 0.00165698 |
| ethanolamine | 0.002184853 |
| xanthine | 0.002247063 |
| 5-methoxytryptamine | 0.002300687 |
| uracil | 0.002623214 |
| 7490 | 0.002920079 |
| isoleucine | 0.003141742 |
| 100730 | 0.004886865 |
| 110321 | 0.005167197 |
| cysteine | 0.0060314 |
| cholesterol | 0.006123783 |
| 105554 | 0.006963892 |
| taurine | 0.007193771 |
| tryptophan | 0.007265821 |
| 111776 | 0.008882999 |
| 95422 | 0.011249879 |
| inosine | 0.011381472 |
| glyceric acid | 0.011381472 |
| tyrosine | 0.012384678 |
| 120562 | 0.012645068 |
| 104906 | 0.015896638 |
| methionine | 0.019772261 |
| 42003 | 0.022147104 |
| 1029 | 0.026186395 |
| glycerol-alpha-phosphate | 0.028414301 |
| **22423** | 0.028414301 |
| **myristic acid** | 0.030490244 |
| **pseudo uridine** | 0.033967137 |
| **oxoproline** | 0.038094748 |
| **aspartic acid** | 0.038094748 |
| **133244** | 0.039519768 |
| **108936** | 0.045954398 |
| **lactic acid** | 0.052656987 |

**Supplemental Table 5.**
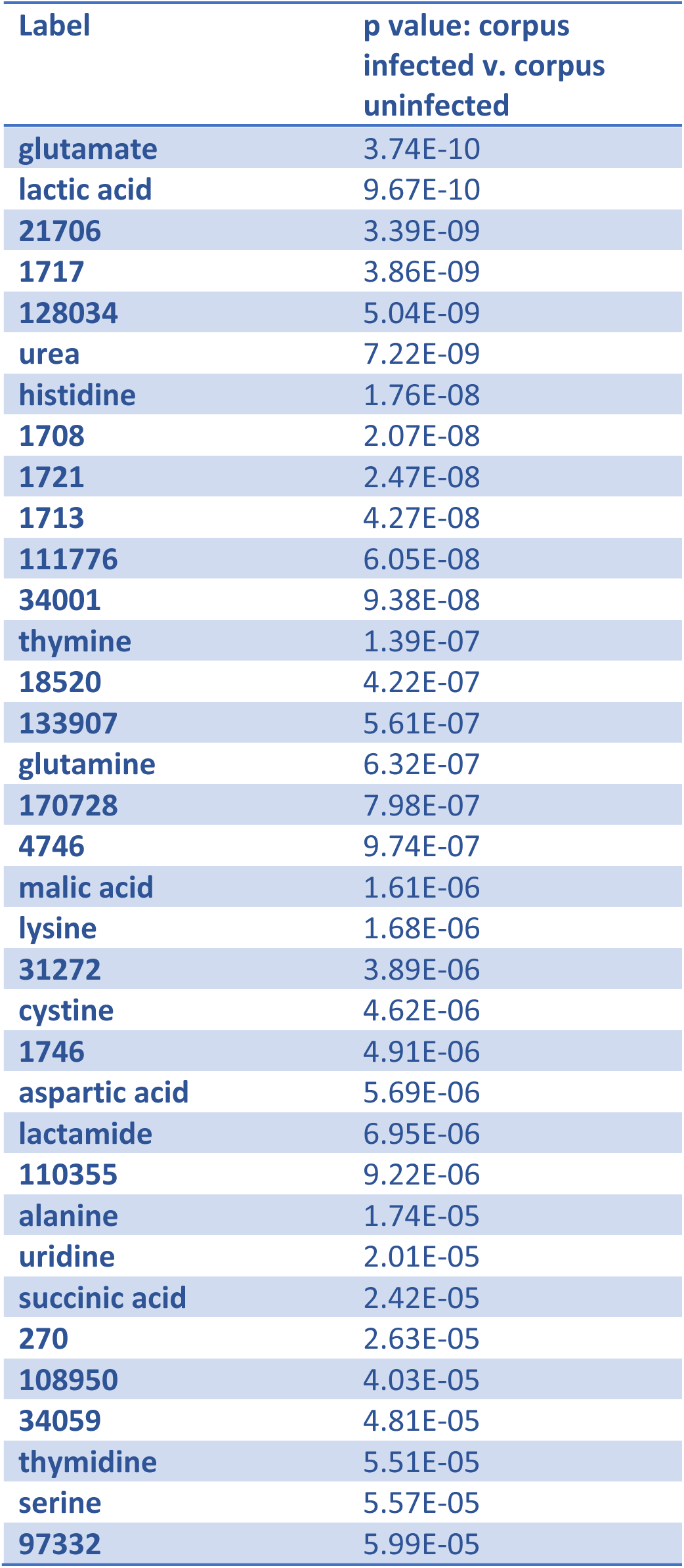
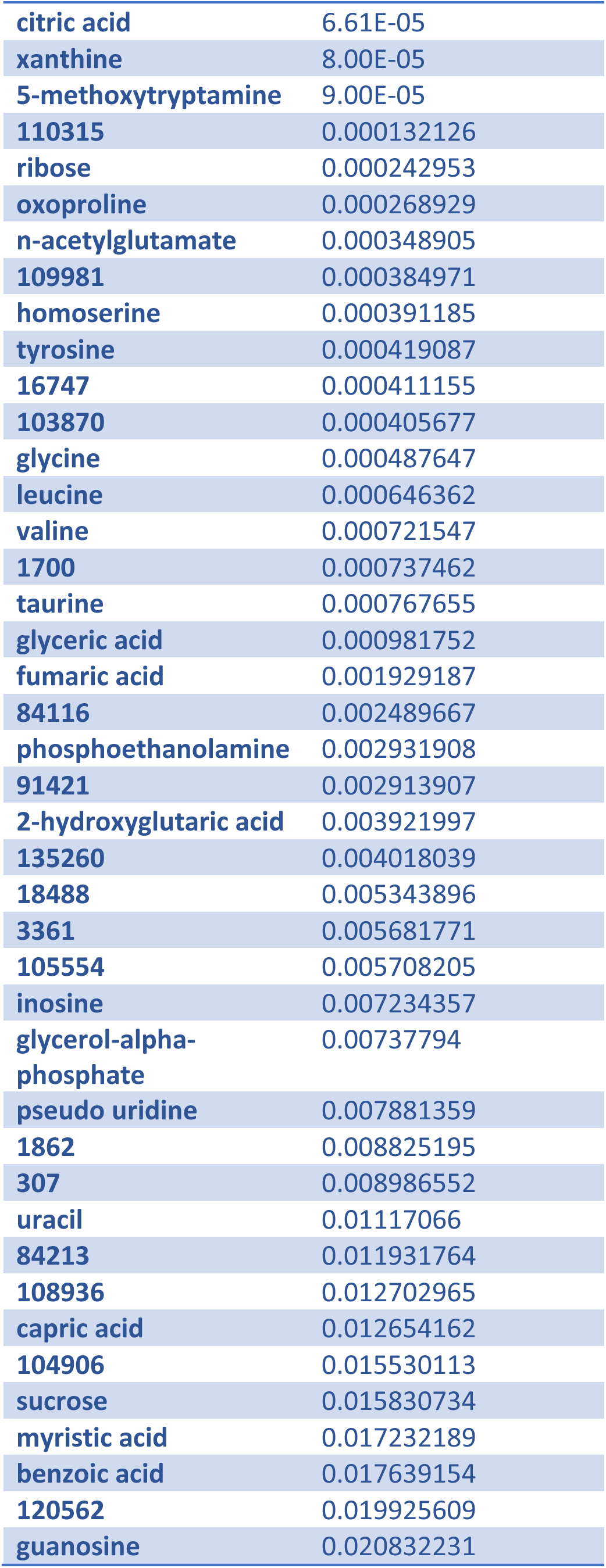
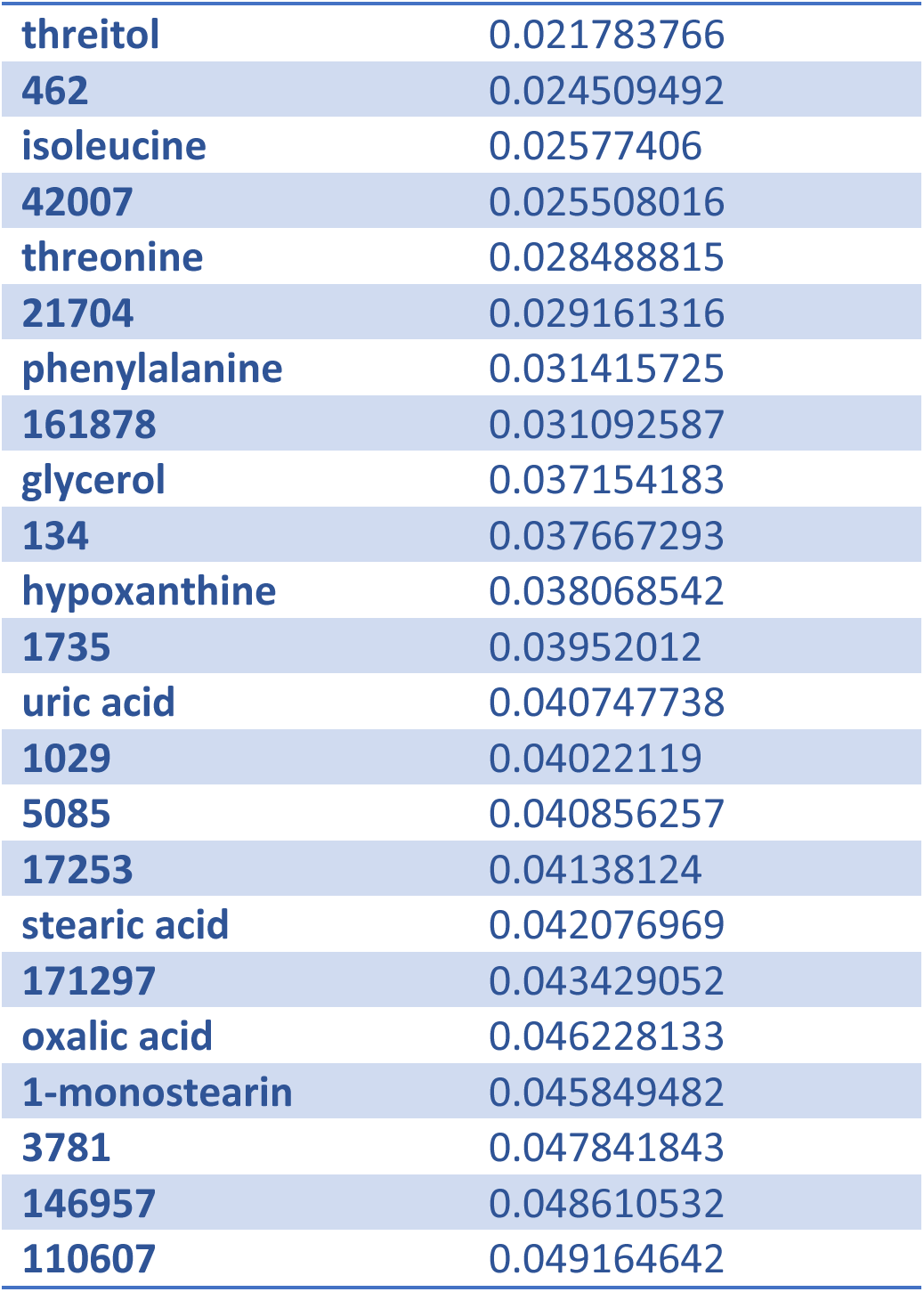
All compounds that changed significantly in the corpus organoids upon infection, ordered by P value.

| Label | p value: corpus<br>infected v. corpus<br>uninfected |
| --- | --- |
| glutamate | 3.74E-10 |
| lactic acid | 9.67E-10 |
| 21706 | 3.39E-09 |
| 1717 | 3.86E-09 |
| 128034 | 5.04E-09 |
| urea | 7.22E-09 |
| histidine | 1.76E-08 |
| 1708 | 2.07E-08 |
| 1721 | 2.47E-08 |
| 1713 | 4.27E-08 |
| 111776 | 6.05E-08 |
| 34001 | 9.38E-08 |
| thymine | 1.39E-07 |
| 18520 | 4.22E-07 |
| 133907 | 5.61E-07 |
| glutamine | 6.32E-07 |
| 170728 | 7.98E-07 |
| 4746 | 9.74E-07 |
| malic acid | 1.61E-06 |
| lysine | 1.68E-06 |
| 31272 | 3.89E-06 |
| cystine | 4.62E-06 |
| 1746 | 4.91E-06 |
| aspartic acid | 5.69E-06 |
| lactamide | 6.95E-06 |
| 110355 | 9.22E-06 |
| alanine | 1.74E-05 |
| uridine | 2.01E-05 |
| succinic acid | 2.42E-05 |
| 270 | 2.63E-05 |
| 108950 | 4.03E-05 |
| 34059 | 4.81E-05 |
| thymidine | 5.51E-05 |
| serine | 5.57E-05 |
| 97332 | 5.99E-05 |
| citric acid | 6.61E-05 |
| xanthine | 8.00E-05 |
| 5-methoxytryptamine | 9.00E-05 |
| 110315 | 0.000132126 |
| ribose | 0.000242953 |
| oxoproline | 0.000268929 |
| n-acetylglutamate | 0.000348905 |
| 109981 | 0.000384971 |
| homoserine | 0.000391185 |
| tyrosine | 0.000419087 |
| 16747 | 0.000411155 |
| 103870 | 0.000405677 |
| glycine | 0.000487647 |
| leucine | 0.000646362 |
| valine | 0.000721547 |
| 1700 | 0.000737462 |
| taurine | 0.000767655 |
| glyceric acid | 0.000981752 |
| fumaric acid | 0.001929187 |
| 84116 | 0.002489667 |
| phosphoethanolamine | 0.002931908 |
| 91421 | 0.002913907 |
| 2-hydroxyglutaric acid | 0.003921997 |
| 135260 | 0.004018039 |
| 18488 | 0.005343896 |
| 3361 | 0.005681771 |
| 105554 | 0.005708205 |
| inosine | 0.007234357 |
| glycerol-alpha-phosphate | 0.00737794 |
| pseudo uridine | 0.007881359 |
| 1862 | 0.008825195 |
| 307 | 0.008986552 |
| uracil | 0.01117066 |
| 84213 | 0.011931764 |
| 108936 | 0.012702965 |
| capric acid | 0.012654162 |
| 104906 | 0.015530113 |
| sucrose | 0.015830734 |
| myristic acid | 0.017232189 |
| benzoic acid | 0.017639154 |
| 120562 | 0.019925609 |
| guanosine | 0.020832231 |
| threitol | 0.021783766 |
| 462 | 0.024509492 |
| isoleucine | 0.02577406 |
| 42007 | 0.025508016 |
| threonine | 0.028488815 |
| 21704 | 0.029161316 |
| phenylalanine | 0.031415725 |
| 161878 | 0.031092587 |
| glycerol | 0.037154183 |
| 134 | 0.037667293 |
| hypoxanthine | 0.038068542 |
| 1735 | 0.03952012 |
| uric acid | 0.040747738 |
| 1029 | 0.04022119 |
| 5085 | 0.040856257 |
| 17253 | 0.04138124 |
| stearic acid | 0.042076969 |
| 171297 | 0.043429052 |
| oxalic acid | 0.046228133 |
| 1-monostearin | 0.045849482 |
| 3781 | 0.047841843 |
| 146957 | 0.048610532 |
| 110607 | 0.049164642 |

**Supplemental Table 6.**
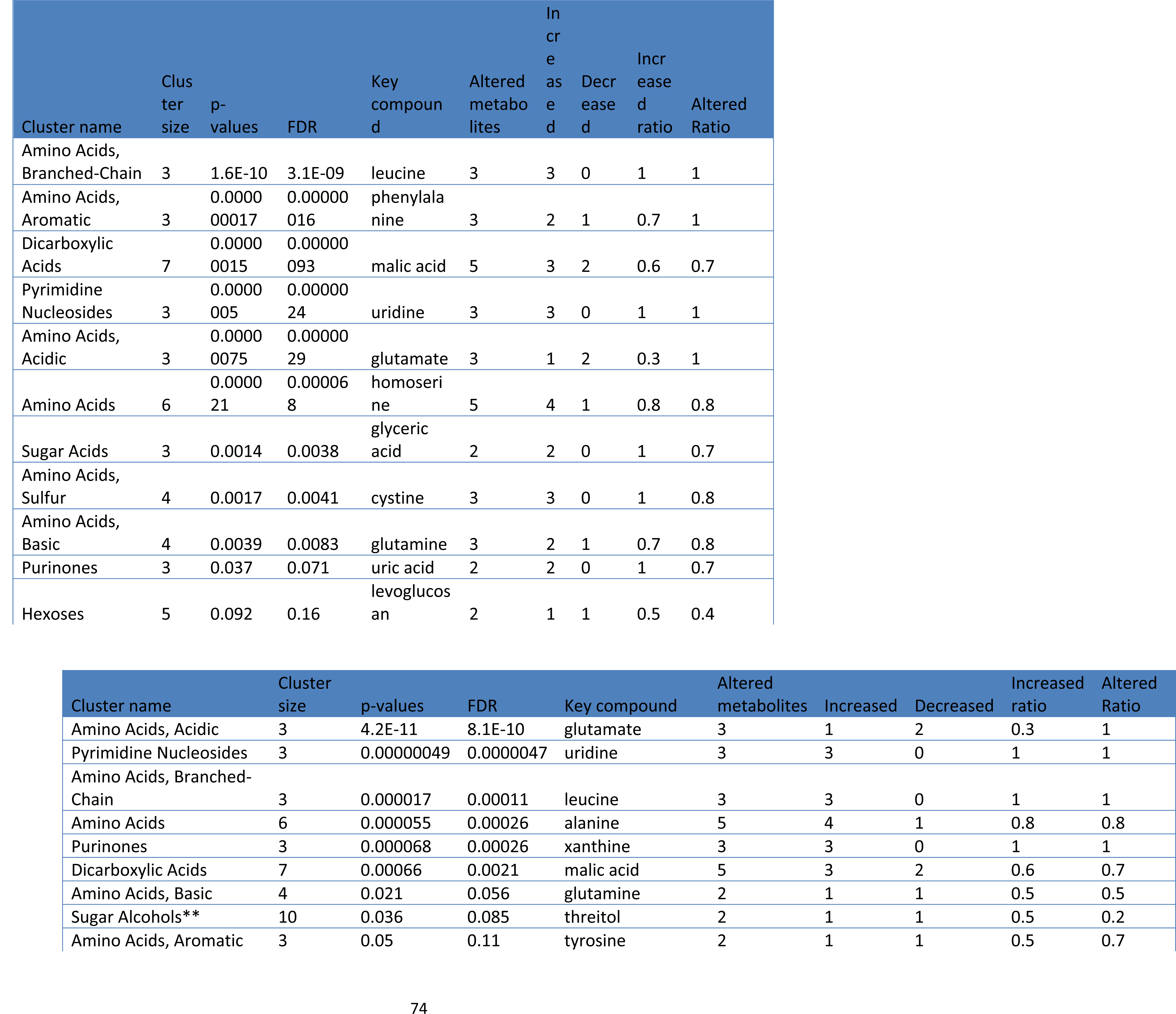
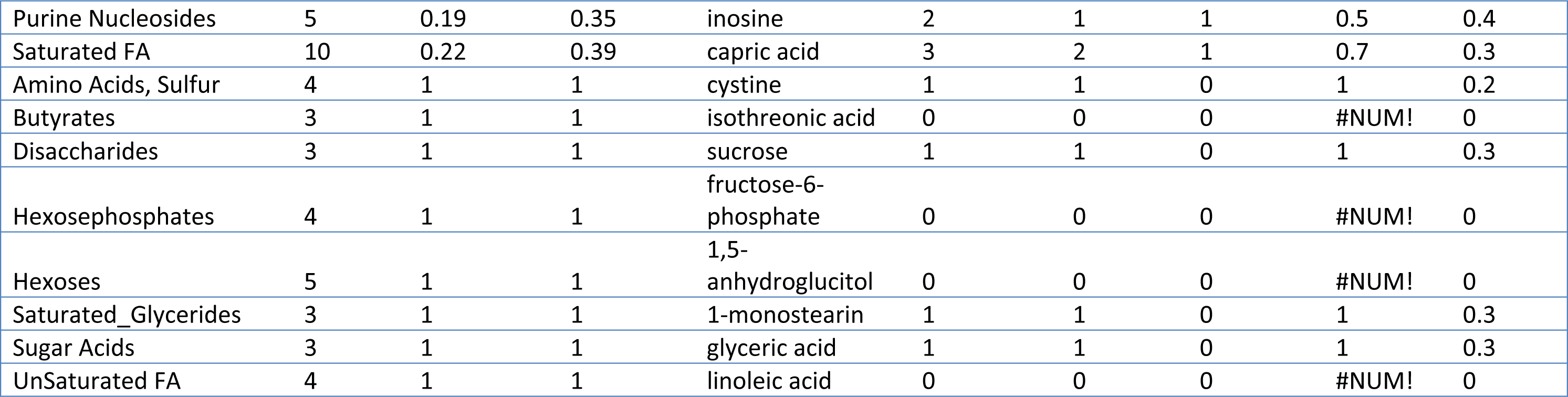
Enrichment analysis comparing uninfected and infected tissues

**Supplemental Table 7.** All chemicals that differed between uninfected antrum and corpus mouse tissue

| Metabolite | p value | FDR corrected p value |
| --- | --- | --- |
| 573 | 0.000 | 0.001 |
| 121402 | 0.000 | 0.001 |
| ethanolamine | 0.000 | 0.001 |
| 97455 | 0.000 | 0.001 |
| 50367 | 0.000 | 0.003 |
| 31357 | 0.000 | 0.003 |
| arachidonic acid | 0.000 | 0.005 |
| 5-methoxytryptamine | 0.000 | 0.006 |
| 1875 | 0.000 | 0.006 |
| linoleic acid | 0.000 | 0.006 |
| 5990 | 0.000 | 0.006 |
| glycerol-alpha-phosphate | 0.000 | 0.006 |
| 111263 | 0.000 | 0.006 |
| oleic acid | 0.000 | 0.007 |
| fucose | 0.001 | 0.008 |
| 31563 | 0.001 | 0.008 |
| mannitol | 0.001 | 0.008 |
| 41808 | 0.001 | 0.008 |
| 2-monopalmitin | 0.001 | 0.011 |
| lactic acid | 0.001 | 0.012 |
| sucrose | 0.001 | 0.012 |
| 135260 | 0.002 | 0.013 |
| uric acid | 0.002 | 0.013 |
| 21683 | 0.002 | 0.018 |
| 84922 | 0.002 | 0.018 |
| 1862 | 0.003 | 0.021 |
| 21706 | 0.004 | 0.027 |
| 443 | 0.004 | 0.029 |
| 100253 | 0.005 | 0.033 |
| 6066 | 0.006 | 0.033 |
| 146957 | 0.006 | 0.033 |
| 2061 | 0.006 | 0.033 |
| octadecanol | 0.006 | 0.033 |
| squalene | 0.006 | 0.033 |
| 16572 | 0.006 | 0.036 |
| dodecanol | 0.008 | 0.044 |
| 2936 | 0.008 | 0.044 |
| 892 | 0.010 | 0.050 |
| 42003 | 0.010 | 0.051 |
| 31285 | 0.011 | 0.051 |
| 106936 | 0.012 | 0.054 |
| 84181 | 0.012 | 0.054 |
| 84383 | 0.012 | 0.054 |
| 1-monoolein | 0.012 | 0.055 |
| 103870 | 0.013 | 0.056 |
| pyruvic acid | 0.013 | 0.056 |
| 62304 | 0.014 | 0.057 |
| 134336 | 0.014 | 0.058 |
| 4901 | 0.015 | 0.060 |
| 102711 | 0.015 | 0.060 |
| fructose-6-phosphate | 0.016 | 0.060 |
| 1029 | 0.016 | 0.060 |
| glutamine | 0.016 | 0.060 |
| glucose | 0.017 | 0.062 |
| 110604 | 0.018 | 0.062 |
| benzoic acid | 0.019 | 0.062 |
| adenosine-5-monophosphate | 0.019 | 0.062 |
| 42007 | 0.020 | 0.062 |
| phosphate | 0.020 | 0.062 |
| 17068 | 0.020 | 0.062 |
| 1700 | 0.020 | 0.062 |
| 1878 | 0.020 | 0.062 |
| 307 | 0.023 | 0.069 |
| 46292 | 0.023 | 0.069 |
| oxalic acid | 0.023 | 0.069 |
| inosine | 0.023 | 0.069 |
| hexose-6-phosphate | 0.024 | 0.070 |
| palmitic acid | 0.025 | 0.071 |
| 17651 | 0.027 | 0.076 |
| threonine | 0.028 | 0.076 |
| 100730 | 0.029 | 0.078 |
| 31359 | 0.029 | 0.079 |
| glucose-6-phosphate | 0.030 | 0.080 |
| capric acid | 0.033 | 0.086 |
| pelargonic acid | 0.034 | 0.088 |
| 1709 | 0.035 | 0.089 |
| **10962** | 0.036 | 0.089 |
| **hydroxycarbamate NIST** | 0.036 | 0.089 |
| **3258** | 0.037 | 0.090 |
| **5346** | 0.038 | 0.091 |
| **1912** | 0.038 | 0.091 |
| **hydroxylamine** | 0.041 | 0.097 |
| **657** | 0.043 | 0.100 |
| **131620** | 0.045 | 0.103 |
| **succinic acid** | 0.046 | 0.105 |
| **malic acid** | 0.048 | 0.106 |
| **caprylic acid** | 0.048 | 0.106 |
| **fumaric acid** | 0.049 | 0.106 |
| **112533** | 0.049 | 0.106 |
| **uridine** | 0.050 | 0.107 |
| **creatinine** | 0.050 | 0.107 |

**Supplemental Table 8.** Compounds that changed between uninfected and 8 month infected corpus mouse tissue

| label | p value: uninfected * 8 months<br>@corpus tissue | adjusted p value: uninfected * 8<br>months @corpus tissue |
| --- | --- | --- |
| threonine | 0.00000878 | 0.000506869 |
| cholesterol | 0.0000154 | 0.000506869 |
| inosine | 0.0000161 | 0.000506869 |
| glycine | 0.000023 | 0.000506869 |
| xanthine | 0.0000235 | 0.000506869 |
| uric acid | 0.000019 | 0.000506869 |
| adenosine | 0.0000105 | 0.000506869 |
| 46128 | 0.00000824 | 0.000506869 |
| squalene | 0.0000201 | 0.000506869 |
| 54 | 0.0000462 | 0.000896776 |
| 257 | 0.0000548 | 0.000967324 |
| aminomalonate | 0.0000853 | 0.001378536 |
| ornithine | 0.0000968 | 0.001444545 |
| urea | 0.000153078 | 0.002121219 |
| 146957 | 0.00020686 | 0.002508179 |
| methionine | 0.00026525 | 0.003026965 |
| aspartic acid | 0.000405264 | 0.004367841 |
| oxoproline | 0.000464323 | 0.004740981 |
| creatinine | 0.00056276 | 0.005458773 |
| 111263 | 0.000726901 | 0.006715183 |
| 112533 | 0.000848875 | 0.006861743 |
| 97455 | 0.000789351 | 0.006861743 |
| glutamine | 0.000847102 | 0.006861743 |
| 573 | 0.001219807 | 0.009465704 |
| 42914 | 0.001275955 | 0.009520586 |
| putrescine | 0.001347462 | 0.009681766 |
| alanine | 0.001534679 | 0.01015553 |
| fucose | 0.001489801 | 0.01015553 |
| 2061 | 0.001570443 | 0.01015553 |
| cysteine | 0.001652174 | 0.01033941 |
| lactic acid | 0.001790385 | 0.01039079 |
| 5990 | 0.001874627 | 0.01039079 |
| 121402 | 0.001821545 | 0.01039079 |
| **50367** | 0.001831048 | 0.01039079 |
| **1872** | 0.002138795 | 0.011388238 |
| **isoleucine** | 0.0022465 | 0.011388238 |
| **1709** | 0.002289388 | 0.011388238 |
| **phosphate** | 0.002252143 | 0.011388238 |
| **palmitic acid** | 0.00261724 | 0.012693614 |
| **arachidic acid** | 0.003124121 | 0.014782426 |
| **serine** | 0.003259147 | 0.015054154 |
| **1-monoolein** | 0.004247252 | 0.019162021 |
| **linoleic acid** | 0.004709791 | 0.020765897 |
| **arachidonic acid** | 0.005676693 | 0.023940836 |
| **47** | 0.005556421 | 0.023940836 |
| **hypoxanthine** | 0.005846673 | 0.024133074 |
| **131620** | 0.005989265 | 0.024206614 |
| **84383** | 0.006173727 | 0.024442917 |
| **oleic acid** | 0.00653424 | 0.02535285 |
| **31359** | 0.006897578 | 0.026237847 |
| **valine** | 0.007432717 | 0.027639269 |
| **171305** | 0.007550934 | 0.027639269 |
| **31357** | 0.008206771 | 0.029483586 |
| **5576** | 0.008573751 | 0.030048426 |
| **hydroxylamine** | 0.008673772 | 0.030048426 |
| **caprylic acid** | 0.009114464 | 0.031021159 |
| **1,5-anhydroglucitol** | 0.009965453 | 0.032767761 |
| **5242** | 0.009934638 | 0.032767761 |
| **stearic acid** | 0.010922961 | 0.035317573 |
| **fructose-6-phosphate** | 0.011504352 | 0.036587612 |

**Supplemental Data Set 1, Organoid Data and Mouse Data from 3d and 28d: MX269377**

**Supplemental Data Set 2. Mouse Data: uninfected and 8 month infected**

